# AID facilitates TET2 demethylation of *Irf4* for plasma cell fate in germinal center B cells

**DOI:** 10.1101/2025.05.02.651162

**Authors:** Minghui He, Roberta D’Aulerio, Lia G. Pinho, Evangelos Doukoumopoulos, Tracer Yong, Rhaissa C. Vieira, Mariana M.S. Oliveira, Rômulo G. Galvani, Saikiran Sedimbi, Christina Seitz, Nikolai V. Kuznetsov, Maria A. Zuriaga, Daisuke Kitamura, José J. Fuster, Vinicius Cotta-de-Almeida, Lena Ström, Søren E. Degn, Lisa S. Westerberg

## Abstract

Activation-induced cytidine deaminase (AID) is essential for B cell affinity maturation. We investigated why AID deficiency gives rise to giant germinal centers (GC) using the AID^R112H^ mouse model that is devoid of AID activity. The increased GC response was associated with accumulation of GC B cells in the light zone in immunized AID^R112H^ mice. AID^R112H^ GC B cells had reduced capacity to up-regulate IRF4 to initiate plasma cell differentiation, leading to accumulation of a transitional GC population with reduced GL7 expression. Genetic introduction of a high affinity B cell receptor (BCR) was unable to restore plasma cell differentiation of AID^R112H^ B cells while ectopic expression of catalytic active AID rescued plasma cell generation. AID^R112H^ impaired recruitment of AID to the *Irf4* promoter/enhancer and disrupted the interaction with Ten-eleven Translocation 2 (TET2). Consequently, DNA demethylation at the *Irf4* promoter/enhancer was reduced in AID^R112H^ GC B cells and impeded high *Irf4* expression for transition into plasma cells. This data reveals a B cell-intrinsic mechanism that governs the plasma cell fate decision through epigenetic remodeling mediated by AID in cooperation with TET2.

**Key messages:**

- AID deficiency leads to accumulation of a transitional (t)GC population of B cells with low expression of GL7 and failure to up-regulate IRF4
- AID deamination activity is needed to promote up-regulation of IRF4 for plasma cell differentiation.
- Co-operation between TET2 and AID facilitates demethylation of the *Irf4* enhancer/promoter.

## Introduction

A hallmark of an efficient humoral immune response is the production of affinity matured antibodies and formation of isotype-switched, long-lived plasma cells and memory B cells in germinal centers (GC). A mature GC can be histologically divided into a dark zone (DZ) and a light zone (LZ). Proliferative B cells populating the DZ highly express activation-induced cytidine deaminase (AID) to diversify their B cell receptors (BCR) through somatic hypermutation (SHM) and Ig class switch recombination (CSR)^1,2^ ^3,4^. The somatically mutated B cells test their BCR affinity through capturing antigens deposited on follicular dendritic cells (FDC) and presentation to T follicular helper (Tfh) cells in the LZ^5^. External signals from FDCs and Tfh cells intertwined with epigenetic and transcriptional programs to orchestrate GC B cells differentiation into plasma cells^6^.

As B cells transit through the GC response to form plasma cells, the genomic architecture is reconfigured substantially^7^. This is mediated through marked epigenetic remodeling including alterations in histone modification and DNA methylation^6^. To counteract the passive DNA demethylation induced by rapid cell proliferation, GC B cells highly express a DNA methyltransferase, DNMT1, to maintain the B cell specific DNA methylome pattern^8^. During DNA replication, DNMT1 recognizes the hemi-methylated CpG site and methylates CpG on the newly synthesized DNA strand^9^. On the other hand, locus-specific DNA demethylation occurs in GC B cells to ensure normal GC B cell differentiation^10^. In mammals, active DNA demethylation is mediated by the members of Ten-eleven Translocation (TET) protein family, including TET1, TET2 and TET3^11^. The TET protein can mediate iterative oxidation of 5mC, therefore generating 5-hydroxymethylcytosine (5hmC), 5-formylcytosine (5fC), and 5-carboxylcytosine (5caC)^12^. 5hmC, 5fC and 5caC impair the DNMT1 binding, which can lead to passive DNA demethylation through replication-dependent dilution of 5mC. Alternatively, 5fC and 5caC can be excised by thymine DNA glycosylase (TDG) that is coupled with base excision repair pathway to mediate active DNA demethylation^11^. Among the three TET proteins, both TET2 and TET3 are expressed in GC B cells^13^. The loci undergoing DNA hypomethylation in GC B cells is overrepresented at B cell enhancers that cover the key genes essential for plasma cell formation, for example *Irf4, Prdm1 and Xbp1*^14^. Concomitantly, up-regulation of IRF4 and down-regulation of BCL6 and PAX5 precede the plasma cell differentiation^15^. Recent work suggests that an intermediate expression of IRF4 is required for the GC response, while high and sustained level of IRF4 is required for plasma cell differentiation^16^. This coincides with a progressive DNA demethylation at the *Irf4* locus from naïve B cells to terminal differentiated plasma cell stage and is mediated by TET proteins^17^. In contrast, differentiation of memory B cells from GC seems to be a default fate for GC B cells as long as they have been selected for survival^18,19^.

To initiate SHM and CSR, AID is associated with the transcription machinery and specifically targeted to the Ig variable and switch regions, where AID can deaminate cytidine into uracil at the single stranded DNA (ssDNA) exposed during transcription^20–24^. The resulting U:G mismatch can be sensed and removed by Uracil-DNA glycosylase (UNG), eliciting a DNA damage response ^25–28^. Repair of AID induced DNA damage will result in SHM in the variable region and CSR in the constant region of the Ig locus^1,2^. AID deficiency leads to a primary immunodeficiency disease called type II hyper-IgM syndrome^29^. A hotspot mutation identified in type II hyper-IgM patients is Arginine 112 to histidine mutation (AID^R112H^). Arginine 112 is located adjacent to the substrate recognition loop in the APOBEC domain. The R112H mutation abolishes the catalytic activity of the AID protein^30–33^. In addition to the complete absence of SHM and CSR, both type II hyper-IgM patients and AID deficient mice possess lymphadenopathy caused by giant and persistent GC^4,29,34^. The cause of the persistent GC as a result of AID deficiency remains to be determined.

In addition to BCR diversification, it has been suggested that AID mediates active demethylation probably via deamination of 5-methylcytosines in various non-B cell models^35,36^. Although AID preferentially deaminates unmethylated cytosines, biochemical evidence from independent studies have demonstrated deamination activity of AID towards 5mC^37,38^. Moreover, marked methylome alteration in AID deficient GC B cells has been identified in mouse models and Hyper IgM syndrome patients^39,40^. In addition, AID drives methylome heterogeneity and accelerates GC-derived B cell lymphomagenesis^41^. However, direct evidence for the relevance of AID-mediated DNA demethylation in GC B cells is lacking.

Using our previously established mouse model for the type II hyper IgM syndrome ^34^, we found that lymphadenopathy in AID^R112H^ mice was caused by accumulation of GC B cells with low expression of GL7 at a transitional GC stage. This was caused by reduced capacity to induce plasma cell differentiation due to compromised upregulation of IRF4 in AID^R112H^ GC B cells. Genetic introduction of a high affinity BCR failed to rescue the plasma cell differentiation in AID^R112H^ GC B cells. Instead, ectopic expression of wildtype AID protein in AID^R112H^ B cells fully restored the capacity to generate plasma cells. Our data reveals that AID and TET2 cooperate at the *Irf4* promoter/enhancer, and that AID deamination activity was required for efficient DNA demethylation of the *Irf4* gene for commitment to the plasma cell lineage.

## Results

### Increased GC response with accumulation of light zone B cells in AID^R112H^ mice

Expression of AID is detectable throughout B cell development with highest expression in GC DZ B cells ^42,43^. We analyzed the possible impact of the AID^R112H^ mutation on early B cell development from the pro/pre-B cells stage in the bone marrow to mature B cell stages in spleen. B cell development in bone marrow and spleen was comparable between AID^R112H^ and wildtype (WT) mice (Supplemental Figure 1A and 1B). To examine the B cell intrinsic effect of the AID^R112H^ mutation and to exclude an effect of the microenvironment in AID^R112H^ mice, we generated mixed bone marrow chimeric mice that were reconstituted with a 1:1 ratio of CD45.1 WT and CD45.2 AID^R112H^ cells. Circulating B cells and marginal zone B cells in the chimeric mice were comparable between WT and AID^R112H^ cells. There was a slight advantage of AID^R112H^ cells in the follicular B cell and transitional B cell populations (Supplemental Figure 1C). This suggested that the antigen-independent phase of B cell development in the bone marrow and spleen was not impaired by the AID^R112H^ mutation.

**Figure 1.**
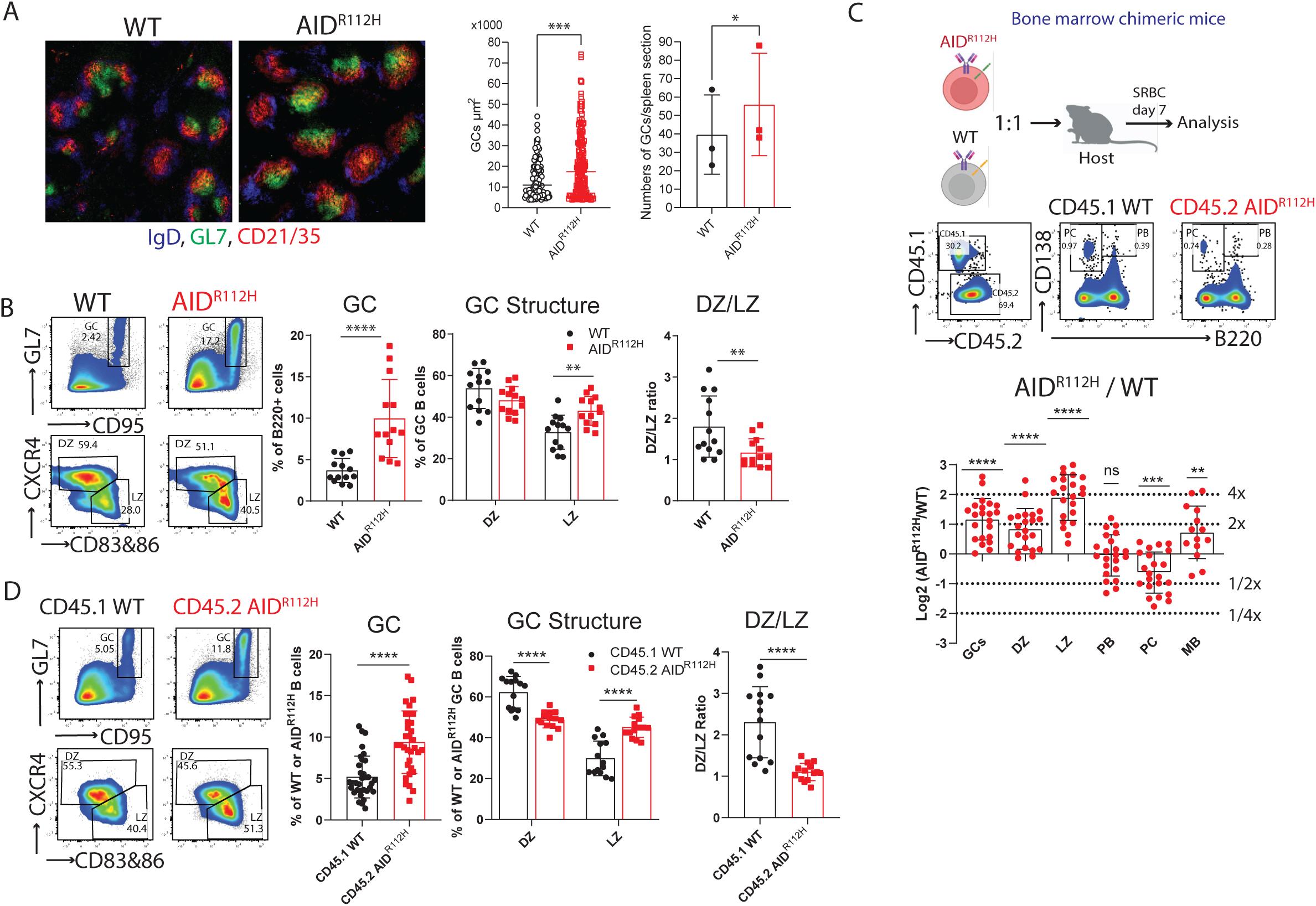
Increased GC response with altered structure in AID^R112H^ mice and bone marrow chimeric mice. (A) GC in spleen section on day 7 upon SRBC immunization. Blue: IgD; green: GL7; red: CD21/35. (B) GC response on day 7 upon SRBC immunization. Left panels: representative FACS plot for GC, DZ and LZ; Right panels: quantification, WT vs AID^R112H^; GC, p<0.0001; DZ, p=0.137; LZ, p=0.0037. DZ/LZ ratio, p=0.0097, unpaired *t* test, (n=13, three experiments). (C) Antigen-dependent B cell differentiation in the spleen of bone marrow chimeric mice. Upper panel: experiment scheme. Middle panel: FACS plot of splenic PB and PCs in the chimeric mice. Lower panel: quantification, y-axis: log2 of measured AID^R112H^/WT to grafted AID^R112H^/WT ratio; GC, DZ and LZ: p<0.0001; Plasma cell, PC: p<0.001; Memory B, MB: p<0.01 one sample t test, (n= 14 to 22, three experiments). (D) GC response in the bone marrow chimeric mice. Left panels: representative FACS plot; Right: quantification, CD45.1 WT vs CD45.2 AID^R112H^; GC, p<0.0001, paired *t* test, (n=33, four experiments); DZ, p<0.0001; LZ, p<0.0001, two-way ANOVA (n=14, three experiments); DZ/LZ ratio, p<0.0001, paired *t* test, (n=14, three experiments). Value and error bar: mean±SD.

Immunohistochemistry and flow cytometry analysis revealed that AID^R112H^ mice showed increased GC response with elevated GC B cells when compared to WT mice (Figure 1A-B)^34^. AID^R112H^ GC B cells had a reduced active caspase 3^+^ population when compared to WT cells (Supplemental Figure 1D), which is consistent with previous findings of reduced apoptosis in AID knockout GC B cells^44,45^. To study the GC response in detail, we analyzed GC B cells (GL7^+^CD95^+^) and among these, LZ (CXCR4^lo^CD83/86^hi^) and DZ (CXCR4^hi^CD83/86^lo^) populations, the generation of spleen plasmablasts (PB, B220^+^CD138^+^), plasma cells (PC, B220^-^CD138^+^), and memory B cells (MB, PD-L2^+^CD73^+^) during the GC response upon SRBC immunization. The GC in AID^R112H^ mice had altered structure, with increased LZ population leading to a reduced DZ to LZ ratio in AID^R112H^ mice when compared to WT mice (Figure 1B). AID^R112H^ mice showed a more than two-fold increase of the memory B cell population (Supplemental Figure 1E). The output of total spleen plasmablasts and plasma cells was comparable between AID^R112H^ and WT mice (Supplemental Figure 1F). In the competitive setting of bone marrow chimeric mice, CD45.2 AID^R112H^ B cells had 2-fold accumulation of GC B cells when compared to CD45.1 WT B cells (Figure 1C and 1D). Moreover, CD45.2 AID^R112H^ GC B cells predominantly located in the LZ whereas the CD45.1 WT GC B cells distributed normally between the two zones (Figure 1D). This demonstrated a cell-intrinsic effect of AID^R112H^ GC B cells even in an environment where the WT cells could sustain a productive GC response and where IgG from WT B cells could mediate antibody feedback regulation^46^. Consistent with the increased memory B cell population in individually immunized AID^R112H^ mice (Supplemental Figure 1E), CD45.2 AID^R112H^ B cells showed an advantage in memory B cell formation in competition with CD45.1 WT B cells (Figure 1C). WT and AID^R112H^ cells contributed equally to differentiation of plasmablasts and there was a disadvantage of plasma cell formation from the AID^R112H^ GC B cells (Figure 1C). These data show that AID deficiency led to aberrant accumulation of GC B cells with preferential localization in the LZ of the GC.

### Unaltered cell proliferation rate but compromised G1/S transition of AID^R112H^ GC B cells

To study if the accumulation of GC B cells in AID^R112H^ mice was caused by increased cell proliferation, SRBC-immunized WT and AID^R112H^ mice were given a 4-hour BrdU pulse before analysis on day 7. DZ B cells contained a higher proportion of BrdU^+^ cells when compared to the LZ cells (Supplemental Figure 2A). Surprisingly, GC B cells from AID^R112H^ mice had a decreased BrdU^+^ cell population when compared to GC B cells from WT mice regardless of their location in DZ or LZ (Supplemental Figure 2A). Cell cycle distribution of GC B cells showed an increased G1 population and a decreased early S phase cell population in AID^R112H^ mice when compared to WT mice (Supplemental Figure 2A). These data suggested a reduced fraction of proliferating B cells in AID^R112H^ GCs compared to WT GCs, which is likely due to compromised G1/S transition of AID^R112H^ GC B cells. This may be caused by an impaired BCR affinity maturation in AID^R112H^ mice, since capacity of GC B cell division is proportional to the amount of captured antigens^47^. To investigate whether there is defective DNA replication/initiation in AID^R112H^ B cells, we set up a system using the induced germinal center B (iGB) cells to separate cells that newly initiated the G1/S transition from those already in the DNA synthesis phase of the cell cycle^48^. iGB cells were first given a 15 minutes EdU pulse, followed by BrdU labeling for 5 min, 1 h, 2 h and 4 h. Cells that were in S phase during both EdU and BrdU labeling were EdU^+^BrdU^+^, while cells that were in G1 phase during EdU labeling and had newly entered S phase during BrdU labeling were EdU^-^BrdU^+^. By measuring the increase of the BrdU single positive cells, we could measure the rate of G1/S transition. AID^R112H^ iGB cells showed a reduced G1/S transition rate when compared to WT iGB cells (Supplemental Figure 2B). Consistently, cell cycle distribution of iGB cells showed increased G1 and decreased S phase cell populations among AID^R112H^ iGB cells when compared to WT iGB cells (Supplemental Figure 2B). Despite decreased G1/S transition of AID^R112H^ cells, the global cell proliferation rate in AID^R112H^ iGB cells was unaffected when compared to WT iGB cells as shown by the dilution of cell division dye Cell Trace Violet (CTV) (Supplemental Figure 2C). Together, these data excluded that the lymphadenopathy and accumulation of GC B cells in AID^R112H^ mice was caused by faster GC B cell proliferation rate.

### AID^R112H^ mice have reduced plasma cell differentiation in GC and accumulates GL7^lo/-^ transitional GC B cells

Our data from bone marrow chimeric mice showed an advantage of AID^R112H^ B cells to form GC B cells but a disadvantage to contribute to the plasma cell pool. We reasoned that the large GCs formed in AID^R112H^ mice were caused by a blockage of plasma cell differentiation from GC B cells. Plasma cells can differentiate from both the GC pathway and the extrafollicular pathway ^15^. To investigate if there was defective plasma cell formation from GC B cells in AID^R112H^ mice, we performed a new gating strategy as shown in Figure 2A to examine the plasma cell differentiation within the GC. Newly formed plasmablasts (PB) and plasma cells (PC) were analyzed in the DZ and LZ of the GL7^+^CD95^+^ GC population (Figure 2A). AID^R112H^ mice had a reduced CD138^+^ population (including both PB and PC) from both DZ and LZ cells when compared to WT mice (Figure 2A). The majority of CD138^+^ cells in the LZ were PB (Figure 2A). In contrast, in the DZ population most of the CD138^+^ cells had down-regulated the cell surface receptor B220 and become PC (Figure 2A). Similarly to individual mice, analysis of bone marrow chimeric mice showed that GC B cells from CD45.2 AID^R112H^ B cells had a reduced CD138^+^ (PB+PC) population when compared to the CD45.1 WT GC B cells (Supplemental Figure 3A). This data suggests that the AID^R112H^ GC B cells had reduced capacity to differentiate into PB and PC (Figure 2A).

**Figure 2.**
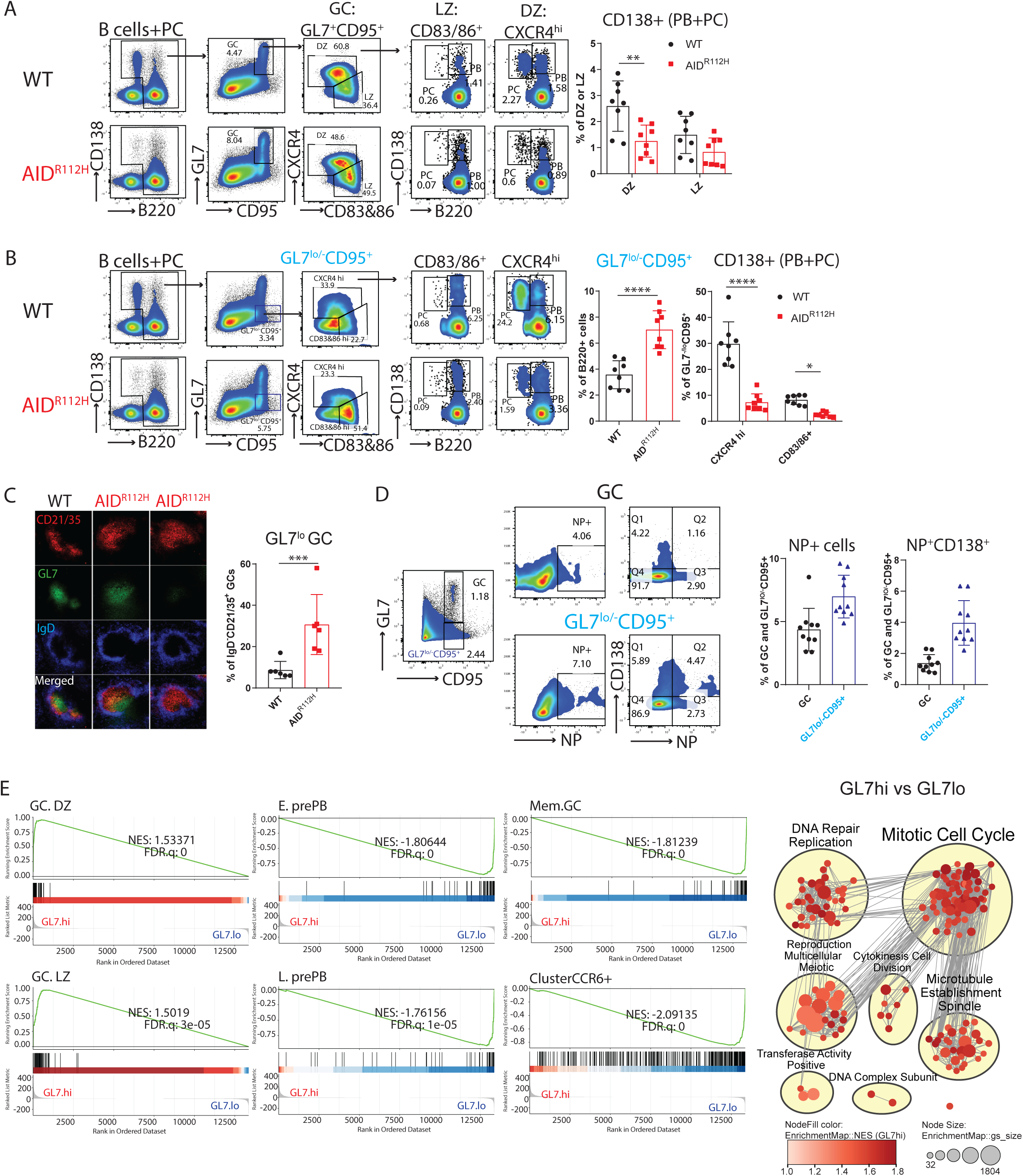
AID^R112H^ mice have reduced plasma cell differentiation in GCs and accumulates GL7^lo/-^ tGC B cells. (A) Gating strategy to identify GC-derived PB and PC. Plasmablast, PB: B220^+^CD138^+^; Plasma cell, PC: B220^-^CD138^+^. FACS plots and quantification, WT vs AID^R112H^; CD138^+^ cells in DZ, p=0.0019; CD138^+^ cells in LZ, non-significant. unpaired *t* test, (n=8, 2 experiments); (B) Gating strategy of PB and PC in GL7^lo/-^CD95^+^ cell population. FACS plots and quantifications, WT vs AID^R112H^; GL7^lo/-^CD95^+^ cells: p<0.0001; CD138^+^ in CXCR4^hi^: p<0.0001; CD138^+^ in CD83/86^+^: p= 0.044. unpaired *t* test, (n=8, 2 experiments). (C) Immunohistochemistry of spleen GCs. Red, CD21/35; Green, GL7; Blue, IgD. Quantification of IgD^-^GL7^-^CD21/35^+^ GCs among total IgD^-^CD21/35^+^ GCs, p=0.0051, unpaired *t* test (n=6, 2 experiments). (D) NP specific response of GC and GL7^lo/-^CD95^+^ B cells in WT mice. FACS plots and quantification, GC vs GL7^lo/-^CD95^+^. NP^+^: p=0.0028; NP^+^CD138^+^: p=0.0002, unpaired *t* test (n=10, 2 experiments). (E) RNA-seq comparing GL7^hi^ vs GL7^lo^ GC B cells. Left panel: Gene set enrichment analysis (GESA) using gene signature defined in work^49,50^; Right panel: GESA network analysis showing gene sets positively enriched in GL7.hi versus GL7.lo. Value and error bar: mean±SD.

In addition to the increased GC B cell population (GL7^+^CD95^+^), AID^R112H^ mice had an increased population of GL7^lo/-^CD95^+^ B cells (Figure 2B). Immunohistochemistry of spleen sections confirmed a significant increase of an aberrant type of GCs in AID^R112H^ mice as marked by IgD^-^CD21/35^+^ area with lower or undetectable level of GL7 compared to WT mice (Figure 2C and Supplemental Figure 3B). Based on the surface expression of CXCR4 and CD83/CD86, these cells could be divided into three populations (Figure 2B; CXCR4^hi^, CD83/86^hi^, and CXCR4^lo^CD83/86^lo^). To examine if this population represented a transitional GC (tGC) B population destined to leave the GC, we analyzed plasma cell differentiation within the GL7^lo/-^ CD95^+^ B cell population. The GL7^lo/-^CD95^+^CXCR4^hi^ B cells contained a significantly higher percentage of CD138^+^ (PB+PC) cells (Figure 2B) when compared to DZ B cells (Figure 2A) in WT mice. This suggested that the GL7^lo/-^CD95^+^ B cells included B cells at a transitional GC stage, and we refer to them as GL7^lo/-^CD95^+^ tGC B cells. In AID^R112H^ mice, the CD138^+^ (PB+PC) cells among the tGC B cells were reduced more than 3-fold when compared to WT mice (Figure 2B). To further validate the transitional GC differentiation stage of the GL7^lo/-^ CD95^+^ B cell population and to be able to identify antigen specific B cells, we immunized WT mice with NP-KLH and examined the GL7^lo/-^CD95^+^ tGC B cells. There was an increased proportion of NP-PE^+^ B cells in the tGC B population when compared to the total GC B cells (Figure 2D). Importantly, there was a more than 2-fold increase of NP^+^CD138^+^ cells in the tGC B cell population when compared to total GC B cells (Figure 2D). In the bone marrow chimeric mice, accumulation of the tGC B population from CD45.2 AID^R112H^ B cells was increased, while generation of CD138^+^ cells in tGC B cells was greatly compromised when compared to CD45.1 WT B cells (Supplemental Figure 3C). Altogether, these results suggest that we have identified a transitional GC population enriched for CD138^+^ plasmablasts and plasma cells. Moreover, plasma cell differentiation from the GC response was significantly compromised in the AID^R112H^ mice and characterized by accumulation of the GL7^lo/-^CD95^+^ tGC B cells with a GC LZ phenotype.

To define the transcriptome profile of tGC B cells, we sorted the GL7^lo/-^CD95^+^ cell population (GL7.lo) and compared to sorted GL7^+^CD95^+^ GC cells (GL7.hi) by RNA sequencing. The GL7^lo/-^CD95^+^ tGC B cells clustered together irrespective of genotype using an unsupervised hierarchical clustering (Supplemental Figure 4A). Gene set enrichment analysis (GSEA) revealed a significant enrichment for signatures of GC DZ and LZ cells in the GL7^hi^ population. In contrast, the GL7^lo/-^CD95^+^ tGC B cell population was significantly enriched for gene signatures of early and late pre-plasmablasts (E.prePB and L.prePB) as well as GC memory B cells (Figure 2E)^49^. Interestingly, GL7^lo/-^CD95^+^ tGC B cells were enriched for a gene signature of CCR6^+^ activated precursor B cells that are a heterogenous population destinated to various cell fates preceding the GC response at day 3-4 upon immunization (Figure 2E)^50^. This suggested that the GL7^lo/-^CD95^+^ B cell was a heterogenous population consisting of precursor cells programmed for further differentiation from both the GC response and extrafollicular pathways. Additionally, the GSEA network analysis revealed a significant enrichment of genes related to cell cycle, cell division and DNA repair/replication in the GL7^+^CD95^+^ (GL7.hi) B cell population, while these gene hallmarks were diminished in the GL7^lo/-^CD95^+^ B cell population (Figure 2E and Supplemental Figure 4B). This suggested a down-regulation of mitotic signatures in the GL7^lo/-^CD95^+^ B cell population. Instead, the GL7^lo/-^CD95^+^ B cells had enriched signatures for cell activation and migration, suggesting an ongoing selection and differentiation process occurring in the LZ (Supplemental Figure 4B). This data further supported the notion that a proportion of the GL7^lo/-^CD95^+^ B cells represented a transitional stage between clonal expansion and terminal differentiation in GCs.

Taken together, the AID^R112H^ GC B cells differentiated into a transitional GC phenotype with reduced GL7 expression and these tGC B cells were halted at the pre-plasmablast stage and failed to differentiate into plasma cells.

### AID^R112H^ GC B cells have reduced capacity to up-regulate IRF4

To investigate the molecular mechanism underlying the defective plasma cell differentiation of AID^R112H^ B cells, we examined the expression of transcription factors BCL6, IRF4, and PAX5 in GC B cells from SRBC immunized mice. Around 90% of the GL7^+^ GC B cells were BCL6^+^ in both WT and AID^R112H^ mice (Figure 3A). The BCL6**^+^** cells among the GL7^lo^CD95^+^ population was two-fold higher in AID^R112H^ mice compared to WT mice (Figure 3A). This supported the notion that the BCL6^+^ cells accumulated in the GL7^lo^CD95^+^ population in the AID^R112H^ mice. Additionally, AID^R112H^ GCs showed more than 80% reduction of the IRF4^hi^PAX5^lo^ population when compared to WT GCs (Figure 3A). Consistently, the IRF4^hi^PAX5^lo^ population among the tGC B cells was significantly reduced in AID^R112H^ mice when compared to WT mice (Figure 3A). Immunohistochemistry staining of spleen sections from SRBC immunized mice confirmed the presence of cells that highly expressed nuclear IRF4 within the GC area (Supplemental Figure 4C), indicating the initiation of plasma cell differentiation from GC B cells. In addition, we found that there was a significant reduction of IRF4 protein in AID^R112H^ GC B cells when compared to WT cells, while PAX5 protein expression was comparable between AID^R112H^ and WT GC B cells (Figure 3B).

**Figure 3.**
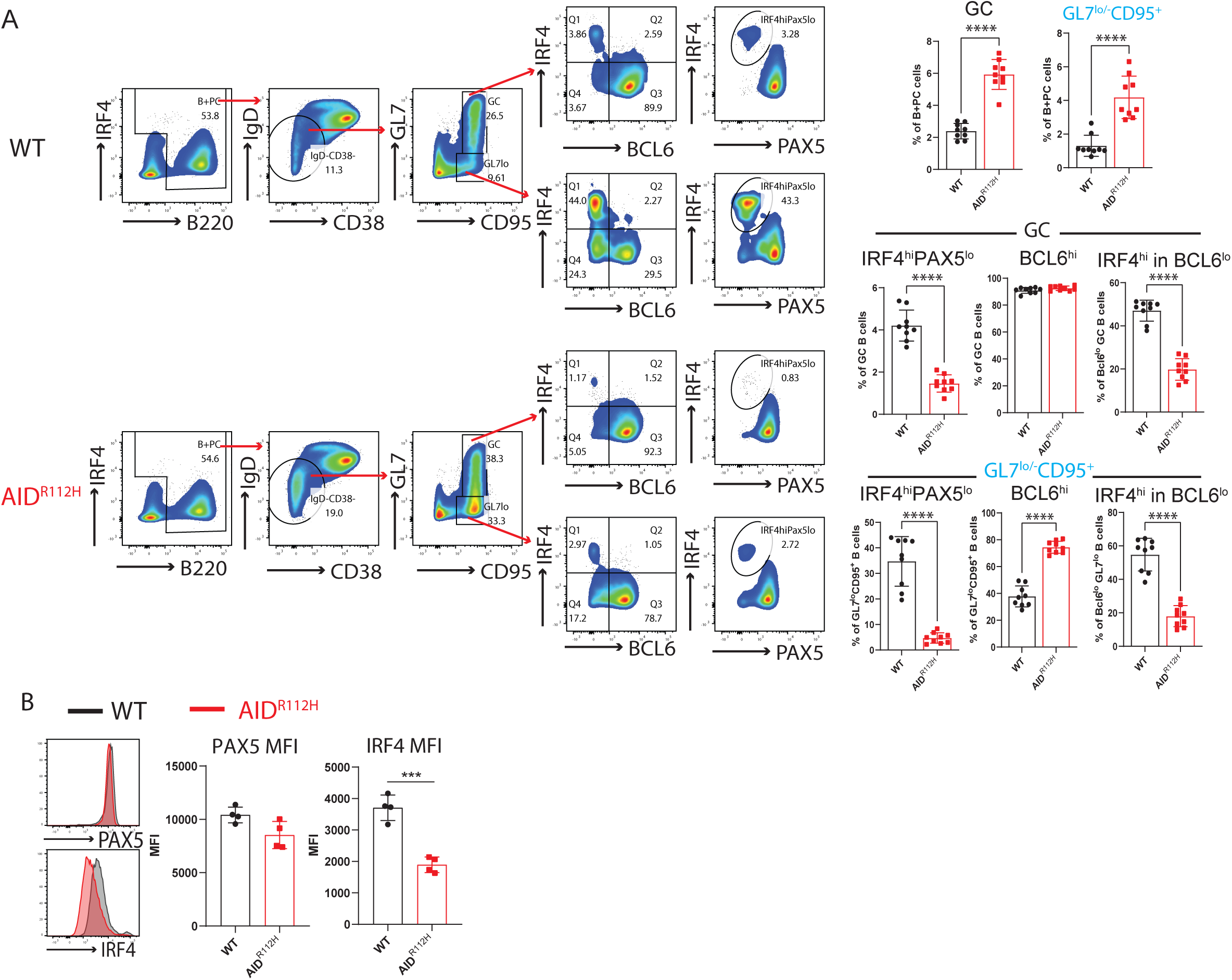
AID^R112H^ GC B cells have reduced capacity to up-regulate IRF4. (A) Gating strategy for expression of transcription factors BCL6, IRF4 and PAX5 in GC and GL7^lo^CD95^+^ B cells. FACS plots and quantification, WT vs AID^R112H^, GC: p<0.0001; GL7^lo/-^CD95^+^, p<0.0001; IRF4^hi^PAX5^lo^ in GC: p<0.0001; IRF4^hi^PAX5^lo^ in GL7^lo/-^CD95^+^, p<0.0001; IRF4^hi^ in BCL6^lo^ (GC): p<0.0001; BCL6^hi^ in GL7^lo^CD95^+^: p<0.0001; IRF4^hi^ in BCL6^lo^ (GL7^lo^CD95^+^): p<0.0001; unpaired *t* test (n=9-10, 3 experiments). (B) Expression level of IRF4 and PAX5 in GC B cells. Overlay of histogram of IRF4 and PAX5 acquired by flow cytometry and quantification, WT vs AID^R112H^, IRF4 MFI: p=0.0006, unpaired t test, (n= 4, 2 experiments). Non-significant differences for PAX5 MFI. Value and error bar: mean±SD. Value and error bar: mean±SD

Previous studies suggest that down-regulation of BCL6 preceded plasma cell differentiation ^51^. GC B cells from AID^R112H^ mice had decreased capacity to down-regulate BCL6 when compared to WT GC B cells (Figure 3A). Moreover, we found that IRF4^hi^ cells only existed in the BCL6^lo^ population. Quantification of the IRF4^hi^ population among the BCL6^lo^ cells showed a signification reduction in AID^R112H^ GCs when compared to WT GCs (Figure 3A). By treating the SRBC-immunized mice with BI-3802, a small molecule inhibitor towards BCL6, we detected an increased BCL6^lo^ population in AID^R112H^ GC B cells. However, increased BCL6^lo^ cells in AID^R112H^ GCs did not lead to increased IRF4^hi^PAX5^lo^ cells (Supplemental Figure 5A). This suggests that down-regulation of BCL6 was not sufficient for IRF4 up-regulation and plasma cell differentiation in AID^R112H^ GC B cells. Moreover, AID^R112H^ B cells had reduced capacity to down-regulate CXCR5 on GC and tGC B cells, providing an explanation for the failure of AID^R112H^ B cells to egress from the GC (Supplemental Figure 5B). These results suggest that AID^R112H^ B cells down-regulated BCL6 and PAX5, but showed reduced capacity to up-regulate IRF4 that is required for GC B cells to differentiate into plasma cells.

### Reduced AID^R112H^ plasma cell formation from *in vitro* induced GC B cells (iGB cells)

To investigate whether the defective plasma cell differentiation in AID^R112H^ mice is B cell intrinsic, we examined the plasma cell differentiation using the iGB cell culture system (Figure 4A) ^48^. Similarly to plasma cells generated in vivo, B cells differentiating into plasmablasts down-regulated PAX5 and up-regulated IRF4 and CD138 (Figure 4A). On day 7, a substantial proportion of cells had differentiated into plasmablasts in the WT cell culture (Figure 4B and 4C). Consistent with our *in vivo* data, the differentiation of plasmablasts from AID^R112H^ B cells was significantly compromised (Figure 4B and 4C). Moreover, iGB cells derived from AID^R112H^ B cells showed reduced amount of IRF4 protein when compared to WT iGB cells, while PAX5 protein was similar in WT and AID^R112H^ iGB cells (Figure 4D). Moreover, the impaired up-regulation of IRF4 in AID^R112H^ iGB cells was not due to compromised cell proliferation (Supplemental Figure 5C). Analysis of iGB cells using imaging flow cytometry confirmed that AID^R112H^ B cells showed greatly reduced nuclear IRF4 when compared to WT B cells (Figure 4E and Supplemental Figure 5D). At the same time, PAX5 and FOXO1, two transcription factors that are present in the GC B cells and down-regulated in plasma cells ^52–55^, showed no difference in iGB cells between WT and AID^R112H^ cells, and indeed were down-regulated in the WT IRF4^hi^PAX5^lo^ population (Supplemental Figure 5D).

**Figure 4.**
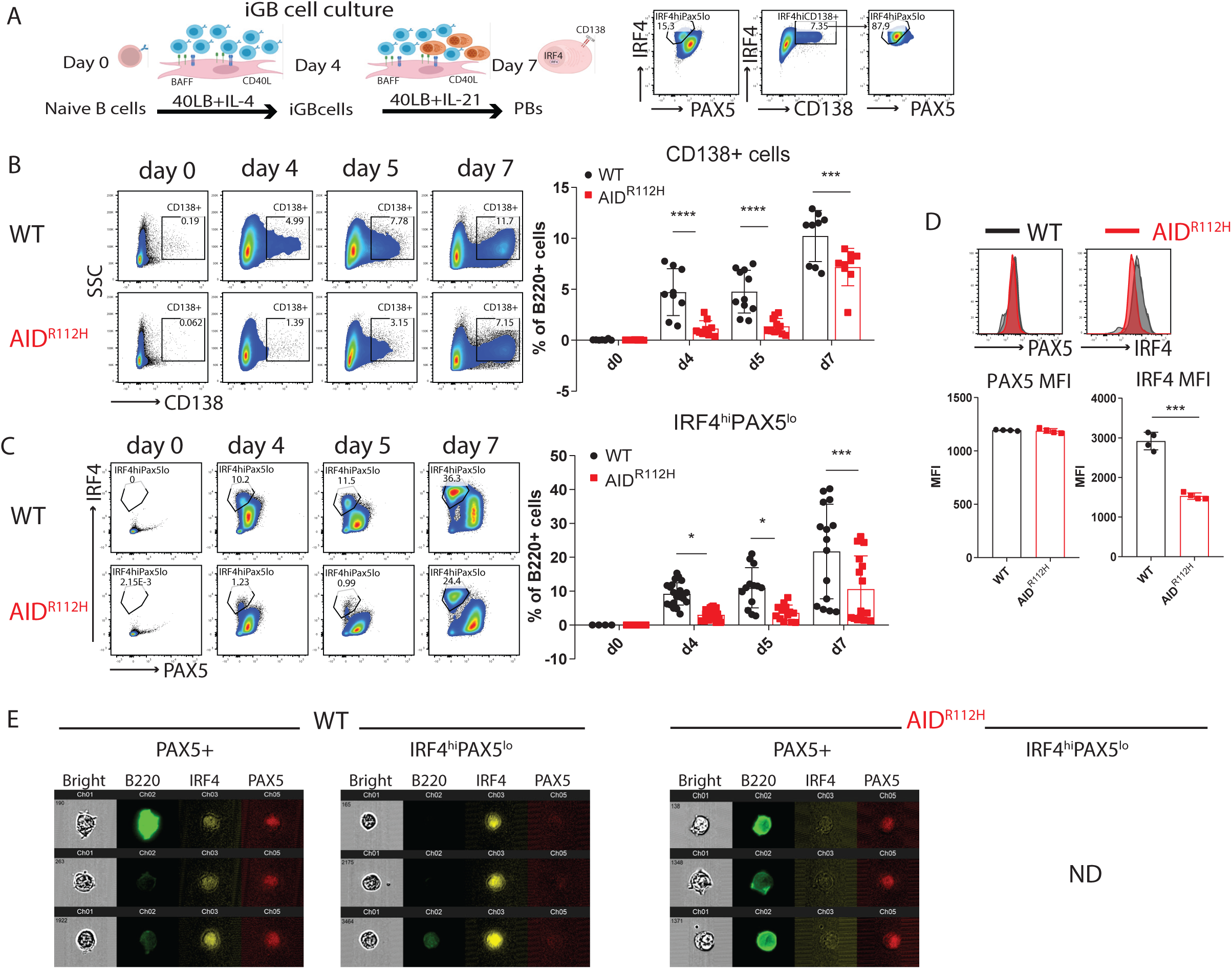
Reduced AID^R112H^ plasma cell formation from *in vitro* induced GC B cells (iGB cells). (A) Plasma cells differentiation in iGB cultures. Schematics of B cells cultured on 40LB cells and representative FACS plots for plasmablast differentiation (CD138+ and IRF4^hi^PAX5^lo^ cell population). (B) CD138^+^ PB cells from day 0 to day 7. FACS plots and Quantification. Day 4 (p<0.0001), day 5 (p<0.0001) and day 7 (p=0.0009) (n=9 to 11, 4 experiments). (C) IRF4^hi^PAX5^lo^ cells from day 0 to day 7. FACS plots and Quantification Day 4 (p=0.016), day 5 (p=0.033) and day 7 (p=0.0001) (n=13 to 22, 5 experiments). (D) Expression of IRF4 and PAX5 of iGB cells. Overlay of histograms of IRF4 and PAX5 acquired by flow cytometry. Quantification, IRF4 MFI: p=0.00082, Non-significant difference for PAX5 MFI, unpaired *t* test, (n=4, one representative experiment). (E) Imagestream analysis of IRF4 and PAX5 in iGB cells on day 7. Representative images of PAX5^+^ and IRF4^hi^PAX5^lo^ cells from the iGB culture. Images were acquired with Imagestream flow cytometry (10000 cells acquired, 2 experiments). ND (non-detectable; no cells could be detected). Value and error bar: mean±SD.

We next examined if ectopic expression of IRF4 could rescue defective plasmablast generation in AID^R112H^ iGB cells. Overexpression of IRF4 in the AID^R112H^ iGB cells by retroviral transfection was able to fully restore the IRF4^hi^PAX5^lo^ cell population (Supplemental Figure 5E), showing that up-regulation of IRF4 plays a critical role in promoting plasma cell differentiation. These results validated our *in vivo* findings and showed that the reduced plasmablast generation in AID^R112H^ mice was caused by a B cell-intrinsic defect to up-regulate IRF4.

### Restored AID activity rescues plasma cell differentiation

To examine if the defective plasma cell differentiation of AID^R112H^ GC B cells was caused by compromised affinity maturation, we took advantage of the B1-8^hi^ mice carrying a pre-assembled Ig heavy chain that when paired with an Ig lambda chain produce a BCR with high affinity for the hapten 4-hydroxy-3-nitrophenylacetyl (NP). B1-8^hi^ mice were bred onto AID^R112H^ background (B1-8^hi^AID^R112H^). To compare the NP high affinity WT and AID^R112H^ B cells side by side, we adoptively transferred Cell Trace Violet (CTV)-labeled naïve B cells from B1-8^hi^AID^R112H^ and B1-8^hi^WT mice mixed with 1:1 ratio and analyzed at day 7 upon immunization with NP-CGG (Figure 5A). B1-8^hi^AID^R112H^ and B1-8^hi^WT B cells were activated and underwent comparable cell proliferation as evidenced by the CTV dilution. IRF4^hi^ plasmablasts appeared after 7 divisions from both B1-8^hi^AID^R112H^ and B1-8^hi^WT B cells in vivo (Figure 5A). There was a more than 2-fold reduction of IRF4^hi^ plasmablasts from B1-8^hi^AID^R112H^ B cells as compared to B1-8^hi^WT B cells (Figure 5A). Moreover, there was more than 3-fold reduction of antigen specific plasma cells (IRF4^hi^NP^+^) generated from B1-8^hi^AID^R112H^ cells compared to B1-8^hi^WT B cells (Supplemental Figure 6A), suggesting an impaired high affinity antibody response. Analysis of the GC response revealed that the GC B cell population was composed of both host B cells and adoptively transferred B1-8^hi^AID^R112H^ and B1-8^hi^WT B cells. Although the total transferred B1-8^hi^AID^R112H^ and B1-8^hi^WT B cells maintained a 1:1 ratio 7 days after immunization, there was a more than 2-fold accumulation of B1-8^hi^AID^R112H^ B cells over B1-8^hi^WT B cells in the GC (Figure 5A). Interestingly, we observed a higher percentage of NP^+^ cells among the B1-8^hi^AID^R112H^ GC B population when compared to the NP^+^ cells among the B1-8^hi^WT GC B cells (94.8% vs 65.2%, Supplemental Figure 6B). This led to an increased MFI of surface NP-PE binding of B1-8^hi^AID^R112H^ GC B cells compared to the B1-8^hi^WT cells. A fraction of IgM-expressing cells contained BCR that had lost NP binding in B1-8^hi^WT GC B cells, which was not observed in the B1-8^hi^AID^R112H^ GC B cells (14.2% vs 2.3%, Supplemental Figure 6B). This can be attributed to SHM mediated by AID in the WT cells. Additionally, SHM can also lead to destroyed BCR in GC, resulting in higher fraction of NP^-^ cells in the B1-8^hi^WT compartment. This data together suggests that the defective up-regulation of IRF4 and plasma cell differentiation of AID^R112H^ B cells did not result from impaired affinity maturation.

**Figure 5.**
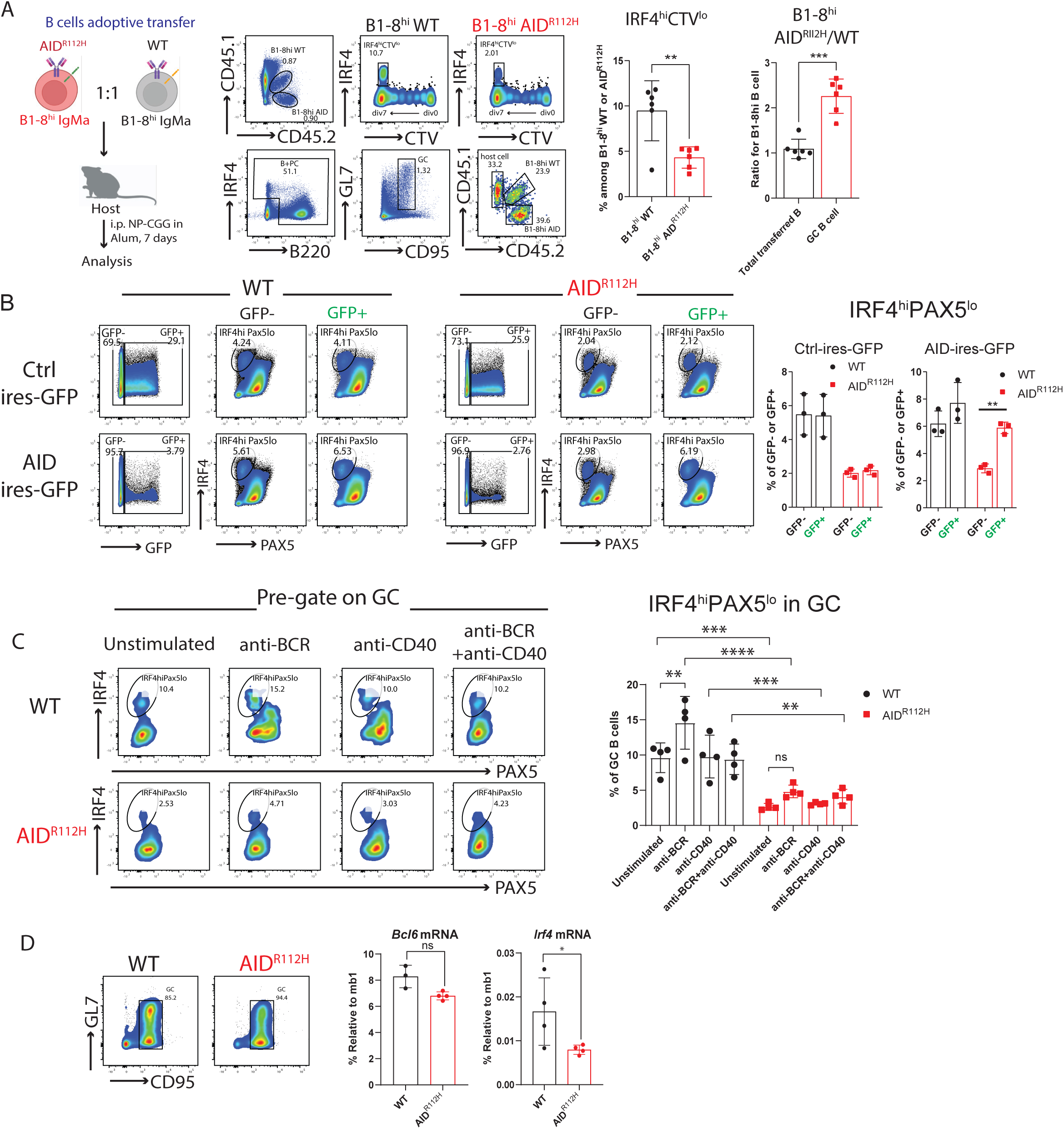
Plasma cell differentiation requires wildtype AID protein. (A) Plasma cell differentiation of B1-8^hi^WT and B1-8^hi^AID^R112H^ cells in vivo. Left: experiment scheme. Middle: Gating strategy to analyze GC response and plasma cell differentiation from adoptively transferred B1-8^hi^WT and B1-8^hi^AID^R112H^ cells; Right: quantification, IRF4^hi^CTV^lo^ in the adoptive transferred WT and AID^R112H^ cells: p=0.005; Ratio B1-8^hi^ AID^R112H^/ B1-8^hi^ WT in GC: p=0.0009; paired t test (n=6, 2 experiments). (B) Rescue of plasmablast differentiation by expressing wildtype AID protein in AID^R112H^ iGB cells. FACS plots and quantification. Ctrl-ires-GFP: no changes between GFP+ vs GFP-in WT and AID^R112H^ iGB cells; AID-ires-GFP: non-significant between WT GFP-vs WT GFP+; p=0.0085 between AID^R112H^ GFP-vs AID^R112H^ GFP+, two-way ANOVA (one representative experiment from three). (C) Percentage of IRF4^hi^PAX5^lo^ cells in GC at day 7 and upon stimulation for 2 hrs. Left: representative FACS plots (pre-gated on GC B cells). Right: quantification, WT, anti-BCR vs unstimulated, p<0.01, non-significant for other stimulations; AID^R112H^, non-significant for all stimulated vs unstimulated comparisons. Two-way ANOVA (n=4 per condition, 2 experiments). (D) *Bcl6* and *Irf4* mRNA level in purified GC B cells. Upper panels: GC purity by flow cytometry. Lower panels: quantification of *Bcl6* and *Irf4* mRNA level by RT-PCR, each dot represents an average value from an individual animal (technical triplicates for each animal), *Bcl6* mRNA, WT vs AID^R112H^, non-significant; *Irf4* mRNA, WT vs AID^R112H^, p<0.05, ratio paired *t* test (n=3-4, 2 experiments). Value and error bar: mean±SD.

To further validate this, we adoptively transferred naïve B cells from B1-8^hi^AID^R112H^ (IgMa, CD45.2) and AID^R112H^ (IgMb, CD45.2) mice, mixed at 1:1 ratio and injected into WT recipient mice (IgMb, CD45.1, Supplemental Figure 6A). At day 7 upon NP-CGG immunization, the plasma cell differentiation from B1-8^hi^AID^R112H^ (IgMa, CD45.2) GC B cells was comparable to the differentiation from AID^R112H^ (IgMb, CD45.2) GC B cells, and both were significantly lower than that from WT (IgMb, CD45.1) GC B cells (Supplemental Figure 7A and 7B).

To rule out that the defective PC differentiation is caused by compromised Ig class switching of AID^R112H^ GC B cells, we analyzed the PC output rate in non-switched (IgM^+^) and switched (IgM^-^) GC B cells from SRBC immunized mice. Indeed, there was comparable level of plasma cell differentiation between IgM^-^ and IgM^+^ GC B cells in wildtype mice, while the IgM^+^ GC B cells from AID^R112H^ mice had decreased capacity to form plasma cells as compared to IgM^+^ B cells from WT mice (Supplemental Figure 8A). Additionally, the capacity of BCR signaling was unaltered in AID^R112H^ GC B cells when compared to WT GC B cells as assessed by p-Syk, p-Btk and p-Erk (Supplemental Figure 8B). Flow cytometry analysis showed comparable level of c-MYC expression in AID^R112H^ GC B cells compared to WT cells (Supplemental Figure 8C-D). Upon stimulation, AID^R112H^ GC B cells up-regulated c-MYC to comparable level of WT GC B cells (Supplemental Figure 8E). These data demonstrate that the defective up-regulation of IRF4 and plasma cell differentiation of AID^R112H^ GC B cells did not result from impaired affinity maturation, Ig class switching and BCR induced up-regulation of c-MYC, indicating that there is another layer of IRF4 regulation mediated by AID activity.

To investigate whether AID activity is directly involved in orchestrating the plasma cell differentiation besides its role in affinity maturation during the GC response, we took advantage of the iGB culture system supplemented with IL-21, which provides crucial signals to support plasma cell differentiation and bypasses the BCR affinity-mediated selection. We expressed wildtype AID protein in the AID^R112H^ iGB cells by retroviral transduction. AID^R112H^ cells that received wildtype AID protein completely restored the IRF4^hi^PAX5^lo^ cell population when compared to AID^R112H^ cells that received a control plasmid (Figure 5B). This establishes that wildtype AID activity plays a critical role in the plasma cell fate decision in addition to its role in affinity maturation-driven selection.

To mimic the molecular consequence of AID deamination activity, we used 5-fluoruracil (5-FU) and pemetrexed (PEM) that elevate the level of deoxyuridine triphosphate (dUTP) and leads to incorporation of deoxyuridine into DNA^56^. 5-FU and PEM treated WT iGB cells had no change in generation of the IRF4^hi^PAX5^lo^ cell population when compared to the non-treated WT cells. In contrast, the AID^R112H^ iGB cells treated with 5-FU and PEM had increased IRF4^hi^PAX5^lo^ populations when compared to non-treated AID^R112H^ cells (Supplemental Figure 9A). This prompted us to investigate whether Uracil DNA glycosylase (UNG) that senses and removes uracil from DNA, is involved in regulation of plasma cell differentiation. We ectopically expressed Ugi, a small peptide inhibitor, in WT iGB cells to inhibit UNG activity^57^. Reduced Ig CSR in Ugi-transduced cells suggested efficient inhibition of UNG activity, while the cell proliferation rate was unaffected in the Ugi-transduced cells compared to the control plasmid-transduced iGB cells (Supplemental Figure 9B-C). WT cells that received the Ugi reduced the IRF4^hi^PAX5^lo^ cell population by more than 50% when compared to cells that received a control plasmid (Supplemental Figure 9D). This data suggests that the AID deamination activity and downstream signals elicited by uracil in DNA are directly involved in orchestrating the plasma cell differentiation during the GC response.

Recent work has demonstrated that the level of IRF4 protein was restricted post-transcriptionally by ubiquitin ligases Cbl and Cbl-b in GC B cells^58^. To understand whether the failure to up-regulate IRF4 in AID^R112H^ GC B cells occurred at the transcriptional level or at the post-transcriptional level, we stimulated WT and AID^R112H^ GC B cells in vitro and examined the IRF4^hi^Pax5^lo^ population. WT GC B cells up-regulated IRF4^hi^PAX5^lo^ cells upon BCR activation, whereas AID^R112H^ GC B cells failed to up-regulate the IRF4^hi^PAX5^lo^ population (Figure 5C). The overall level of the IRF4^hi^PAX5^lo^ population among AID^R112H^ GC B cells was reduced compared to WT GC B cells in both unstimulated and stimulated groups, suggesting a lower mRNA level of *Irf4* in AID^R112H^ GC B cells. We next examined the *Irf4* mRNA level from enriched GC B cells by quantitative (q)PCR. The mRNA level of *Irf4* from AID^R112H^ GC B cells showed a 2-fold reduction, in contrast to the *Bcl6* mRNA that remained comparable between AID^R112H^ and WT GC B cells (Figure 5D). Together, these data suggest that the deamination activity of AID is directly involved in regulating plasma cell fate via up-regulation of *Irf4* mRNA in the GC B cells.

### AID accelerates demethylation of the *Irf4* enhancer/promoter via co-operation with TET2 in GC B cells

Upregulation of IRF4 is associated with a progressive DNA demethylation at the *Irf4* enhancer/promoter region as B cells transit through the GC to form plasma cells^17^. Recent studies have demonstrated a role of AID in regulating the DNA methylome in GC B cells^39,40^. To test whether the impaired *Irf4* transcription was due to compromised DNA demethylation in AID^R112H^ GC B cells, we analyzed the DNA methylation pattern at the *Irf4* enhancer/promoter region (−1471 to −14 bps relative to the first exon) using bisulfite sequencing. This indeed revealed an impaired DNA demethylation at the *Irf4* enhancer/promoter in AID^R112H^ iGB cells when compared to WT iGB cells on day 7 (Figure 6A and Supplemental Figure 9E). TET proteins play a critical role in mediating demethylation of the *Irf4* enhancer/promoter during B cell differentiation into plasma cells^17^. Intriguingly, overall transcriptome and methylome alterations occurring in TET2 knockout GC B cells are largely shared by AID knockout GC B cells^59^. Therefore, we reasoned that AID and TET2 may cooperate to mediate *Irf4* demethylation. Indeed, TET2 knockout (TET2 KO) iGB cells showed a reduction of IRF4^hi^PAX5^lo^ cells compared to WT iGB cells on day 6 (Figure 6B). To examine whether there is a physical interaction between AID and TET2 proteins in B cells, we performed co-immunoprecipitation using iGB cells from day 3 and day 6 cultures. We could detect AID protein from TET2 immunoprecipitations in WT iGB cells but not in the AID^R112H^ iGB cells, suggesting an interaction between AID and TET2 dependent on amino acid R112 (Figure 6C and Supplemental Figure 10A). In contrast, AID was undetectable from the DNMT1 immunoprecipitations from either WT or AID^R112H^ iGB cells (Supplemental Figure 10B). To test if AID was recruited to the *Irf4* enhancer/promoter, we performed Cut&Run sequencing and qPCR using iGB cells. Results from the Cut&Run-sequencing revealed a preferential association of AID to the promoter region of genes (Supplemental Figure 11A). Association of AID to chromatin at global level was reduced in AID^R112H^ iGB cells compared to WT cells as confirmed using two different AID antibodies (Supplemental Figure 11A). Specifically, the association of AID to the known off-target genes involved in the GC response (*Bcl6*, *Pax5*, *Pim1*) and plasma cell differentiation (*Irf4*, *Xbp1*) was diminished in the AID^R112H^ iGB cells compared to WT cells. Association of AID to B cell activation gene *Cd83* was comparable between AID^R112H^ and WT cells, and no association of AID to the *Prdm1* locus was detected (Supplemental Figure 11B-C). Interestingly, the reduced AID-chromatin association was accompanied by a compromised TET2 chromatin association both at the global level as well as at the selected genes (*Bcl6*, *Pax5*, *Pim1, Irf4*, *Xbp1*) in the AID^R112H^ iGB cells (Supplemental Figure 11A-C). In contrast, the global level of H3K4me3 spanning the TSS (transcription start site) had a moderate increase in the AID^R112H^ iGB cells compared to WT cells (Supplemental Figure 11A). To further validate this, we performed qPCR using Cut&Run samples at the selected regions spanning the *Irf4* enhancer/promoter. The results showed a reduced association of AID protein to the *Irf4* enhancer/promoter in AID^R112H^ cells when compared to WT iGB cells on day 3 (Figure 6D). This was accompanied by the impaired demethylation of *Irf4* (Figure 6A) although TET2 was present at the *Irf4* enhancer/promoter in AID^R112H^ iGB cells (Figure 6D). On the other hand, wildtype AID was associated with the *Irf4* enhancer/promoter in TET2 KO iGB cells (Figure 6E). This suggested that AID binding to the *Irf4* locus was independent of TET2. Association of AID to Switch mu (Smu) regions was comparable between AID^R112H^ and WT iGB cells, suggesting differential recruitment mechanisms of AID to non-Ig loci versus Ig loci. Interestingly, although there was a moderate increase of active histone mark H3K4me3 at the global level in AID^R112H^ iGB cells when compared to WT iGB cells, the H3K4me3 at the *Irf4* enhancer/promoter region showed a reduction in AID^R112H^ cells, indicating a reduced *Irf4* transcription initiation/elongation (Supplemental Figure 11A and Figure 6D). This data demonstrates that TET2 and AID cooperate at the *Irf4* promoter/enhancer and that absence of either TET2 or AID activity interferes with de-methylation of the *Irf4* gene to induce transcription.

**Figure 6.**
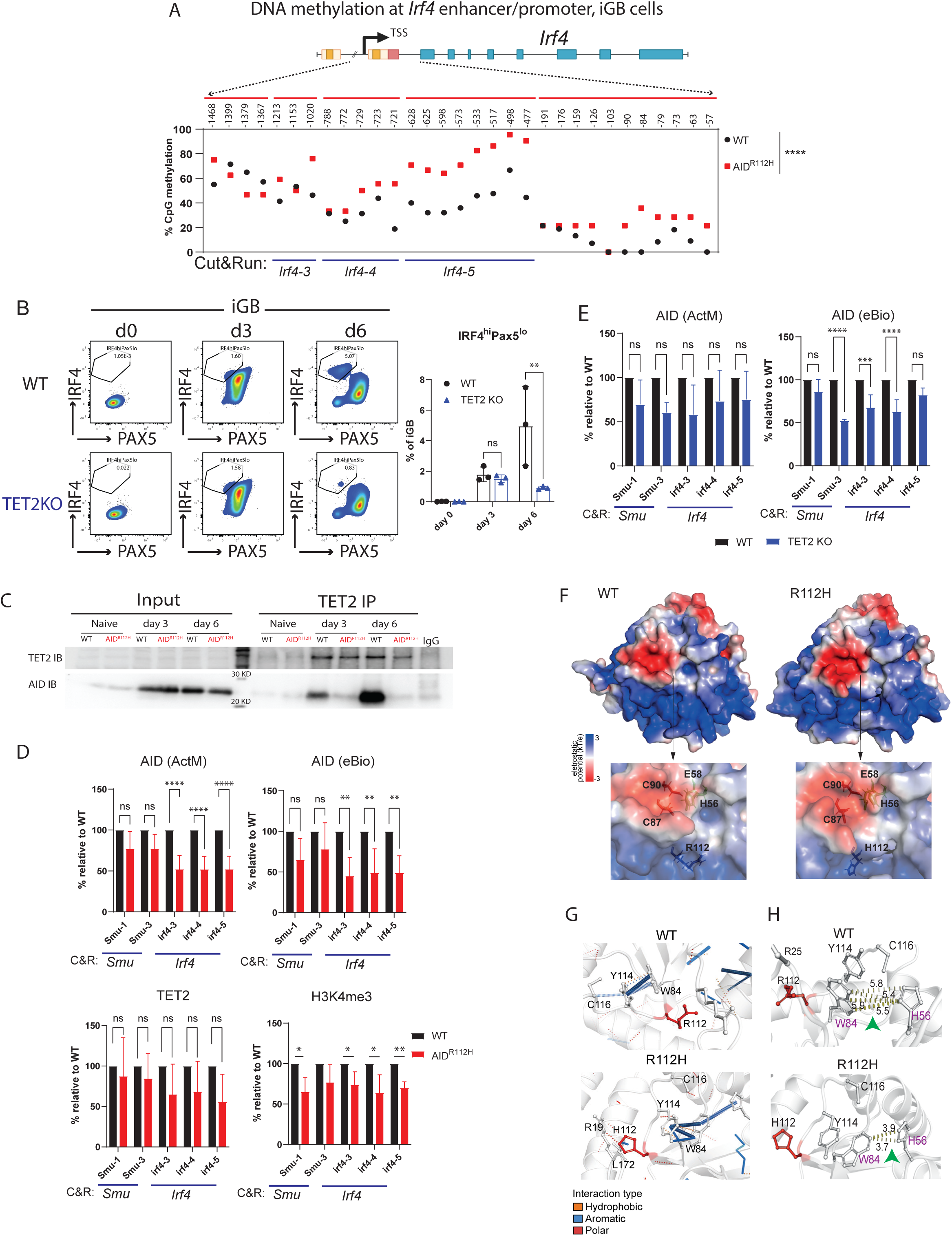
AID co-operates with TET2 to demethylate *Irf4* enhancer/promoter in GC B cells. (A) CpG DNA methylation status at *Irf4* locus in iGB cells derived from WT and AID^R112H^ cells on day 7. Top: schematics for *Irf4* gene locus and the region examined for DNA methylation using bisulfite sequencing. DNA methylation level at CpG sites, number above the graph shows the CpG position relative to the first exon. For each CpG site, 15-30 clones were analyzed, WT vs AID^R112H^, p<0.0001, paired *t* test, data from 2 independent experiments. (B) IRF4^hi^PAX5^lo^ cell differentiation in TET2 KO iGB cells. Representative FACS for day 0, day 3, and day 6. Quantification, day 6, WT vs TET2 KO, p<0.01, two-way ANOVA (n=3, one experiment). (C) Co-IP of endogenous AID and TET2 protein from iGB cells. Anti-TET2 pull down and immunoblot to detect TET2 and AID. One representative blot from 4 independent experiments. (D) Cut&Run and q-PCR to detect association of AID, TET2 and H3K4me3 with the *Irf4* locus in iGB cells on day 3. Cut&Run detection region *Irf4-3*, *Irf4-4* and *Irf4-5* as indicated in the bottom of (A). Black column: WT; Red column: AID^R112H^; Y-axis: % relative to WT, * p<0.05, ** p<0.01, anti-AID (Active Motif and eBioscience), anti-TET2, anti-DNMT1 and anti-H3K4me3. Two-way ANOVA (n=4, four experiments). (E) Cut&Run and Q-PCR to detect association of AID to the Switch mu region and the *Irf4* enhancer/promoter in TET2 KO iGB cells on day 3. Black column: WT; Blue column: TET2 KO; Y-axis: % relative to WT, *** p<0.001, **** p<0.0001, Two-way ANOVA (n=3, three experiments). (F) Electrostatic surface potentials of wildtype AID and AID^R112H^. Upper: whole protein; Lower: catalytic pocket is zoomed in. Blue: positively charged residues; White: neutral residues; Red: negatively charged residues. Position of primary catalytic residues H56, E58, C87, C90 together with R112 are highlighted. (G) Interatomic interactions of affected residues in wild type AID and AID^R112H^. (H) Interaction distance between W84 and H56 in wild type AID and AID^R112H^. Amino acid W84 and H56 are highlighted in purple. Green arrows highlight the interaction distance between the two highlighted amino acids. Wildtype: 5.4-5.9Å; AID^R112H^: 3.7-3.9Å

To further validate AID association and deamination at the *Irf4* locus, we examined the AID-dependent mutational footprint at a GC-rich region of the *Irf4* locus in Ugi-transfected iGB cells (Supplemental Figure 12A). The mutational rate at all three tested regions (Irf4-L1, Irf4-L6 and Irf4-L7) was at the range of 10^-4^ per nucleotide per generation. This is higher than the background genomic mutational rate (10^-6^ per nucleotide per generation), but lower than the estimated somatic hypermutation rate at the IgH locus mediated by AID (10^-3^ per nucleotide per generation). The overall high mutational rate even in AID^R112H^ cells could be attributed to the increased genomic stress induced by retroviral transfection and viral integration into the genome. The mutational rate at the *Irf4*-L1 and *Irf4*-L7 regions was reduced by two-fold in AID^R112H^ iGB when compared to WT iGB cells, while the mutational rate was comparable at the *Irf4*-L6 region between AID^R112H^ and WT iGB cells (Supplemental Figure 12A). This result indicated a deamination activity footprint of AID at the *Irf4* locus. Together, these data demonstrate that AID co-operates with TET2 to facilitate efficient demethylation of the *Irf4* locus to initiate the plasma cell differentiation program.

### In silico structure modelling of AID reveals how R112H abolishes the deaminase activity

To examine the impact of the R112H mutation on protein structure and deamination activity of AID, we performed in silico modelling analysis. The AID RoseTTAFold structures were used to predict ΔΔG of selected mutations (Supplemental Figure 12B-D). Among 26 mutations in AID, the R112H mutation showed an intermediate score (−1.81 kcal/mol) in contrast to the strongly destabilizing mutations, C87S (−4.09 kcal/mol), W80R (−3.69 kcal/mol) and M139T (−3.05 kcal/mol) (Supplemental Figure 12B). The low RMSD values after the alignment of the structures of C87S (0.425 Å), W80R (0.370 Å), M139T (0.372 Å), R112H (0.441 Å), indicated minimal structural difference from the WT AID structure (Supplemental Figure 12C). The H112 residue in AID^R112H^ protein indirectly altered the positive charge near the H56 and the negative charged surface around the catalytic pocket cytosines (Figure 6F). The R112H mutation induced a polar interaction to R19 and an aromatic interaction to L172, leading to reduced interaction distance between W84 and H56 (Figure 6G-H). This resulted in less accessibility to the catalytic pocket of mutant AID^R112H^ protein^61^. Y114 had very high contact frequency with the phosphodiester backbone during ssDNA-AID binding^61^ and Y114 interacted with an oxygen of dCMP^60^. The perturbed interaction between W84 with Y114 could indicate a change in the interaction between AID^R112H^ and ssDNA/dCMP. Indeed, molecular docking of AID^R112H^ with the *Irf4*-AGCT motif had a reduced electrostatic interaction energy compared to the WT-*Irf4*-AGCT complex (Supplemental Figure 12E). Together, these data explain how the R112H mutation in the APOBEC like domain of AID leads to a catalytic dead AID with direct impact on demethylation status of the *Irf4* gene.

## Discussion

Generation of high affinity and long-lived plasma cells from GC is essential for humoral immunity ^62,63^. Extensive studies have revealed that helper signals received from FDC and Tfh cells play a crucial role in regulation of positive selection and plasma cell differentiation in GC ^64–68^. However, the self-intrinsic trigger for B cells to make the plasma cell fate decision remains to be determined. Using the AID^R112H^ mouse model that spontaneously develop GC hyperplasia ^34^, we here demonstrate that lack of AID activity halted GC B cell differentiation at a transitional GC and pre-plasmablast stage, which resulted in the accumulation of GL7^lo/-^ tGC B cells. The GL7^lo/-^ tGC B cells in AID^R112H^ mice failed to up-regulate IRF4 irrespective of BCL6 down-regulation and therefore were unable to initiate the plasma cell differentiation program. Expression of a high affinity BCR was unable to restore the halted plasma cell generation from AID^R112H^ GC B cells in mice. Our data showed that AID and TET2 co-operated at the *Irf4* enhancer/promoter, which accelerated the DNA demethylation and promoted *Irf4* transcription. This data reveals a new layer of regulation mediated by AID to orchestrate plasma cell fate decision through epigenetic regulation. Our data identifies an important example where AID “off-target” activity has direct consequence for cell fate decision during B cell differentiation in GC.

The giant GCs are a characteristic feature of hyper IgM syndrome with AID deficiency, AID KO mice, and AID^R112H^ mice (this study)^4,29,34^. Our work in the current study demonstrated that accumulated GC B cells in AID^R112H^ mice were polarized to the LZ with impaired capacity to generate plasma cells. Affinity-based B cell clonal selection and plasma cell fate commitment occurs in the LZ^64–68^. Therefore, the impaired PC differentiation program in AID^R112H^ GCs likely prevented the cells that have already transited to LZ from leaving the GC or re-entering the DZ, therefore leading to LZ polarization. This is also consistent with previous findings using an AID knockout mouse model^69^. In addition to reduced apoptosis, our work revealed that persistent proliferation, accumulation of atypical tGC B cells, and decreased differentiation to PC together contributed to the enlarged GC in AID^R112H^ mice. Failure to upregulate CXCR5 in the tGC B cells likely contributed to the failure of AID^R112H^ GC B cells to egress from the GC. This data provides a detailed explanation for why AID deficiency leads to the formation of giant GC.

The cell fate decisions of B cells in GC critically depend on the interplay between two groups of transcription factors ^15,63^. Down-regulation of BCL6 precedes plasma cell differentiation of GC B cells ^70,71^. Our study confirmed that the IRF4^hi^ pre-plasmablasts/plasmablasts only existed in the BCL6^lo^ population in GC. The AID^R112H^ mice had reduced capacity to down-regulate the BCL6 protein in GC. Among the BCL6^lo/-^ cells, the IRF4^hi^ population was nearly completely abolished. This suggests that down-regulation of BCL6 alone was not sufficient for IRF4 up-regulation and plasma cell differentiation. These data strongly suggests an additional layer of regulation of IRF4 mediated by AID, which is independent of down-regulation of BCL6 ^72^.

Importantly, expression of wildtype AID protein in the AID^R112H^ iGB cells fully restored generation of IRF4^hi^PAX5^lo^ cells. Treatment with 5-FU or pemetrexed, which could increase the incorporation of deoxyuridine into DNA, partly restored the IRF4^hi^PAX5^lo^ cell population in AID^R112H^ iGB cells. Moreover, inhibition of UNG that removes uracil generated by AID using the small peptide inhibitor Ugi significantly decreased the IRF4^hi^PAX5^lo^ cells in WT iGB cultures, suggesting the involvement of UNG in plasma cell fate decision downstream of AID-mediated deamination. UNG deficiency leads to type 5 hyper IgM syndrome that has similar clinical features as the type 2 hyper IgM syndrome by AID deficiency. This includes lymphoid hyperplasia, increase serum IgM and severely decreased serum IgG and IgA in the UNG deficient patients^73^. However, it is interesting to note that UNG KO and Msh2/6 KO mice have normal or decreased GC response^74–77^. It is likely because UNG and Msh2/6 have pleiotropic effects on proliferating B cells in addition to resolving AID-mediated deamination as critical factors of the base-excision repair and mismatch repair pathways. These data suggested that the deamination activity of AID and the signaling cascade elicited by DNA U:G mismatch plays a critical role in facilitating up-regulation of IRF4.

In addition to lack of IRF4^hi^ cells in AID^R112H^ GC B cells and iGB cells, the protein level of IRF4 in all GC B cells and iGB cells was about 2-fold lower in AID^R112H^ cells when compared to wildtype cells. This suggests that lack of AID activity impacts on a more general process that regulates the IRF4 protein level globally. Indeed, our data demonstrates an impaired transcriptional up-regulation of *Irf4* due to compromised enhancer/promoter demethylation when GC B cells lack AID activity, suggesting an essential role of AID activity in facilitating efficient active demethylation of the *Irf4* locus. However, AID^R112H^ mutant B cells are distinct from the IRF4 KO B cells, because AID^R112H^ B cells were able to up-regulate IRF4 to an intermediate level upon activation. On the other hand, complete deletion of *Irf4* in B cells leads to absence of GC formation as IRF4 KO B cells fail to upregulate BCL6^78^, which is also different from the AID^R112H^ mouse model. Deletion of *Prdm1* (Blimp1) in GC B cells using Cγ1-Cre leads to an increase of GC B cells, which resembles the AID^R112H^ GC^79^. Blimp1 KO B cells can still express AID and deletion of Blimp1 leads to malignant transformation into diffuse large B cell lymphoma in mice and human^78^. Together, this suggests that AID activity is more restricted to facilitate the second stage of *Irf4* demethylation that is required for plasma cell formation.

B cells undergo dramatic epigenetic remodeling and genomic rearrangement upon antigen stimulation^7,10^. Differential DNA methylation occurs as B cells transit from the naïve stage to the GC stage, predominantly consisting of DNA hypomethylation^80^. Concomitantly, GC B cells up-regulate DNMT1 and TET proteins (TET2 and TET3) that exert the opposing function of methylation maintenance and active demethylation to ensure the homeostasis of the GC response. Hypomorphic mutations of DNMT1 in B cells impair GC formation^8^. On the other hand, B cell deficiency for TET2 and TET3 leads to an increased GC response, aberrant accumulation of GC LZ B cells, and impaired plasma cell generation, which resembles the GC response from the AID^R112H^ mice^13,81^ (this study). Our data demonstrate a collaboration between AID and TET2 in GC B cells, which provides a molecular mechanism for the phenotypic resemblance and the shared transcriptome and methylome changes observed in AID knockout and TET2 knockout GC B cells^59^.

Although there is still a controversy regarding a direct role of AID in active demethylation, altered DNA demethylation in GC B cells deficient for AID has been demonstrated by several studies in both mouse models and a Hyper IgM syndrome patient^39,40^. AID binding sites are overrepresented in the hypomethylated loci of GC B cells^8^. Nevertheless, independent studies have shown that the purified AID protein can deaminate 5mCs in vitro although with less activity as compared to unmethylated cytosine deamination. An AID mutant (N51A) protein that only deaminates 5mC can still support substantial level of CSR in vivo, suggesting that deamination of 5mC by AID may still be efficient under the physiological conditions ^82,83^. Moreover, to process TET2-mediated 5mC deoxygenation, cells employ a similar spectrum of repair proteins from the base excision repair (BER) pathway as used for processing of AID-mediated cytosine deamination, e.g. TDG and Apurinic/apyrimidinic endonuclease (APE) ^11^. Synergistic effect of AID and TET2 may greatly augment the DNA damage response and facilitate the recruitment of the shared components from the BER pathway, which in turn accelerates the active demethylation of the *Irf4* enhancer/promoter. The analysis of AID-dependent regulation of DNA demethylation in GC B cells requires development of sophisticated molecular tools to distinguish the role of AID in epigenetic regulation from SHM and CSR. Nevertheless, our data links the deamination activity of AID to epigenetic regulation in the context of B cell differentiation.

The *Irf4* gene has been repeatedly identified as one of the AID off-target genes^84–87^. Our AID Cut&Run data using two different AID antibodies suggested that AID preferentially associated with transcription start sites at its off-target genes, where TET2 showed shared occupancy. Interestingly, at the off-target genes involved in the GC response (*Bcl6, Pax5, Pim1)* and plasma cell differentiation *(Irf4, Xbp1)*, the reduced association of AID^R112H^ was always accompanied by decreased TET2 association to these loci. AID is associated with active transcription to target cytosines in ssDNA formed in the transcription bubble^88^. Our *in silico* modelling provided evidence that the R112H mutation affected the catalytic pocket of AID by indirectly altering the positive charge around H56 and the negative surface around the catalytic pocket. This would impact on the catalytic pocket accessibility, ssDNA binding, interaction with dCMP and possibly, the AID^R112H^ occupancy at on- and off-target sites. Our data shows that for efficient demethylation of the *Irf4* gene, AID and TET2 need to cooperate as neither AID binding to the *Irf4* locus in TET2 KO iGB cells nor TET2 binding to *Irf4* in AID^R112H^ iGB cells is enough to fully demethylate the *Irf4* gene for the high expression of plasma cells.

AID off-targets at non-Ig loci are preferentially associated with B cell super enhancers undergoing convergent/divergent transcription^85,89,90^. This predisposes the GC B cells into increased risk of malignant transformation, as chromosome translocation of IgH to proto-oncogenes is a hallmark of GC-derived B cell lymphomas^91^. However, it is unclear whether there is physiological relevance for AID associating with these non-Ig loci. Recently it has been demonstrated that AID-dependent differentially methylated cytosines are enriched in genes with increased somatic hypermutation burden, and associate with AID-dependent ɣH2AX loci in GC B cells^39^. A central question concerns the mechanism that ensures robust somatic hypermutation in GC B cells to generate high affinity plasma cells, but that also prevents the heavily mutated GC B cells from undergoing tumor transformation. Here we established that in addition to SHM and CSR, AID activity is critical to initiate the plasma cell program in GC by promoting IRF4 expression through locus-specific DNA demethylation. A mechanistic linkage of AID activity and downstream plasma cell lineage commitment probably acts as an additional safeguard against adverse mutational effects of high-level SHM, as it would ensure their sequestration into a terminally differentiated and quiescent cell subset.

## Materials and Methods

### Mice and cell transfers

Mice were bred in the animal facility of KM Wallenberg and Department of Microbiology, Tumor and Cell Biology. All animal experiments performed are approved by the Stockholm North Animal Ethics Committee permits N272/14 and 11159-18. To generate mixed bone marrow chimeric mice, 1×10^7^ mixed bone marrow cells in 1:1 ratio of CD45.1 wildtype cells and CD45.2 AID^R112H^ cells were injected intravenously to the lethally irradiated wildtype congenic recipient mice. B1-8^hi^ IgH knock-in mice were purchased from the Jackson Laboratory and crossed with AID^R112H^ mice to generate B1-8^hi^ AID^R112H^ mice (IgMa, CD45.2)^92^. 5×10^6^ naive spleen B cells mixed in 1:1 ratio from B1-8^hi^ AID^R112H^ mice (IgMa, CD45.2) and AID^R112H^ mice (IgMb, CD45.2) were injected i.v. into wildtype recipient mice (IgMb, CD45.1). This was followed by i.p. immunization with 50 ug NP-CGG per mouse for analysis at day 7. Alternatively, B1-8^hi^ IgH knock-in mice were bred with CD45.1 mice to generate B1-8^hi^ WT (IgMa, CD45.1/CD45.2) mice. 5×10^6^ Cell Trace Violet labeled naive spleen B cells mixed in 1:1 ratio from B1-8^hi^ AID^R112H^ mice (IgMa, CD45.2) and B1-8^hi^ WT (IgMa, CD45.1/CD45.2) mice were injected i.v. into wildtype recipient mice (IgMb, CD45.1), followed by NP-CGG immunization for 7 days.

### B cell isolation and iGB cell culture

Mouse splenic B cells were enriched by the negative B cell selection Kit from Stem Cell Technology. B cells were cultured at 1.6×10^4^ cell/ml on top of 80 Gy-irradiated 40LB cells^48^. Cell culture was supplemented with 1-2 ng/ml of mouse IL-4 (PeproTech). On day 4, 2-4×10^3^ cells from this culture were re-plated on freshly irradiated 40LB cells with 10-20 ng/ml of mouse IL-21 (PeproTech) for another 4-6 days to induce plasma cell differentiation. 10 ng/ml of topoisomerase inhibitor etoposide was added on day 4 for 48 or 96 hrs. 500 ng/ml Fluorouracil (5-FU) or 50 ng/ml pemetrexed was added on day 4 for 24 hrs. To induce T-independent activation of B cells in vitro, 4×10^5^ cells/ml of isolated B cells were culture with complete RPMI1640 with 10 ug/ml LPS for up to 4 days.

### EdU and BrdU labeling

iGB cultures were pulsed with 20 uM of EdU for 15 min and washed away, followed by labeling with BrdU for 5 min, 1 hr, 2 hrs or 4 hrs before harvesting the culture. Detection of EdU was performed using the Click-It EdU kit (Thermofisher). BrdU was detected using monoclonal anti-BrdU antibody.

### Immunization and flow cytometry

Wildtype, AID^R112H^, and WT:AID^R112H^ bone marrow chimeric mice were immunized with sheep red blood cells (SRBC) via i.p. injections. Seven days after immunization, single cell suspensions were prepared from spleen, lymph nodes and bone marrow, and labeled with fluorescence conjugated antibodies including B220, CD43, CD24, CD23, CD21, IgM, IgD, IgG1, CD93, CD138, GL7, CD95, CXCR4, CXCR5, CD83, CD86, CD73, PD-L2, CD80, Blimp1, PAX5, IRF4, BCL6, FOXO1, CD45.1, CD45.2, IgMa, IgMb, anti-BrdU. Data was obtained by LSRFortessa flow cytometry (BD) and analyzed with FlowJo software (TreeStar, Ashland, OR).

### Western blot and Immunoprecipitation

iGB cells were washed with cold PBS and then lysed in RIPA buffer (150mM NaCl, 1.0% IGEPAL, 0.5% Sodium deoxycholate, 0.1% SDS, 50mM Tris, pH 8.0) containing protease and phosphatase inhibitors for 1 hour on ice. For immunoprecipitation, 3ug of antibody was added to cell lysate from 5×10^6^ cells per antibody, and then rotated at 4 degree overnight. Immune complexes were pulled down using 10ul Protein A/G magnetic beads (Thermfisher, Cat. 88803) for 3 hours at 4 degree. Protein was eluted with 60ul of 2xSDS by boiling for 5 minutes at 95 degree. Anti-TET2 (Abcam, Cat. ab124297), anti-DNMT1 (Active Motif, Cat. 39204), anti-AID (Active Motif, Cat. 39885), anti-AID (ThermoFisher, eBioscience, Cat. 14-5959-82), anti-H3K4me3 and anti-IgG (present in CST #91931 kit).

### CUT & RUN (Cleavage under targets and release using nuclease)

5×10^5^ cells were processed with CUT&RUN Assay Kit (Cell Signaling Technology, Cat. #91931) according to manufacturer’s instructions. 4 ug of anti-AID (Active Motif, Cat. 39885), anti-TET2 (Abcam, Cat. ab124297), anti-DNMT1 (Active Motif, Cat. #39204), anti-H3K4me3 and anti-IgG (present in CST #91931 kit) antibodies were used in the experiments. CUT & RUN DNA products were purified using QIAquick PCR purification kit (Cat. #28106) and quantified with real-time PCR using SsoAdvanced SYBR Green Supermix (Bio-RAD, Cat. 1725274). Primer sequences in supplemental Table 1.

### CUT&RUN Sequencing Analysis

Paired-end (150 bp) sequencing was performed by Novogene, using the Illumina Novaseq 6000 platform. Raw reads were quality-filtered with Cutadapt 4.9, and mapped to both Mus musculus (mm39) and Saccharomyces cerevisiae (sacCer3, spike-in) genomes with Bowtie2 2.5.4 ^93^ using the following options; end-to-end--very-sensitive --no-mixed --no-discordant. After normalization with the spike-in raw counts and removal of off-target sequences^94^, normalized bedGraph files were generated with bedtools 2.31.0 (4) and converted to bigWig files using UCSC’s bedGraphToBigWig tool^95,96^. Visualization was done in the UCSC genome browser, and TSS-centered heatmaps and profiles were generated with deepTools 3.5.5^97^.

### DNA methylation analysis by bisulfite treatment

Inducible GC B cells from day 7 were subjected to CpG methylation analysis using EZ DNA Methylation-direct Kit (Zymo Research, Cat. D5021). In brief, 1×10^4^ iGB cells were lysed for 20 minutes at 50 degree with M-digestion buffer containing proteinase K. Cell lysates containing DNA were directly subjected to bisulfite conversion for 3.5 hours. Reaction was stopped using desulphonation buffer, and then DNA was purified with Zymo-Spin column. The modified DNA was amplified by PCR. PCR products were cloned into pCR4-TOPO TA vector and sequenced (ThermoFisher, Cat. K457501). Primer sequences in supplemental Table 1.

### Immunofluorescence staining

Spleens from unimmunized and SRBC-immunized mice were imbedded with Tissue-Tek OCT compound (Sakura) for immunofluorescence staining. 8mm thick tissue sections were cut from the imbedded spleen by using cryostat microtome. Sections were fixed in ice-cold acetone for 10 minutes and blocked with 5% fetal calf serum for 1 h. The sections were incubated with primary antibodies over night at 4°C, washed with PBS, and then incubated with secondary antibodies for 45 min.

### Genotyping

Exon 2 from the *aicda* gene was amplified by Taq polymerase with the following primers: 5’-TCTGGCTGCCACGTGGAATTGT-3’ and 5’-TGATCCCGATCTGGACCCCAGC-3’. PCR program: 95°C, 2min, 30 cycles of 95°C for 30 s, 63°C for 30 s, and 72°C for 30 s, and a last step 72°C for 10 min. The PCR product was digested with 1U BssHII at 37 °C overnight. PCR products from wildtype alleles would be cut into two fragments (175- and 84-bp), whereas the AID^R112H^ product would be resistant to the digestion.

### RNAseq Analysis and Gene Set Enrichment Analysis

Total RNA was purified from sorted cells using RNeasy kit #74104 (QIAGEN). Each RNA sample was treated with RNAse-free DNAse kit #79254 (QIAGEN) during the RNA isolation procedure. The concentration of purified total RNA was measured by NanoDrop 2000 (Thermo Scientific). RNA sequencing was performed using the Illumina HiSeq-PE150 Platform at Novogene, Hongkong, China. Reads were aligned to the reference genome of mouse (Mus musculus) assembly December 2011 (GRCm38/mm10).

The analysis was conducted with R 4.2.3. Briefly, raw gene counts were used as inputs of DESeq2 package (1.38.3) for differential gene expression (DGE) analysis. Prefilter conditions include total gene count < 10 and immunoglobulin variable (IgV) genes. Marker genes from each condition were identified by ranking the product of log fold change and log transformed adjusted p value times −1. Identified Marker genes were visualised using ComplexHeatmap package (2.14.0) with z-score of variance-stablising-transformed (vst) gene counts. The gene set enrichment analysis (GSEA) was performed with clusterProfiler package (4.4.4) with full-ranked DGE result and published marker genes for different stage of B cells^49^ as input files of GSEA command. Pathway enrichment analysis was done in two steps. First, the enriched gene ontology terms was identified for each ranked gene list with gseGO() function. Following that, enriched terms was visualised Cytoscape (3.9.1).

### Structural modelling analysis of AID

The 3D models of wild type and mutated AID were built with RoseTTAFold^98^. PyMOL 3.1 was used for visualization of the structures (https://pymol.org/). The root-mean-square deviation (RMSD) was used to evaluate structural divergence among the predicted models. The mutated AID predicted models were aligned with the wild type AID and the RMSD was measured with PyMOL 3.1. To predict mutation effects on protein stability we calculated the ΔΔG for 26 single point mutations in AID using DDMut web server (https://biosig.lab.uq.edu.au/ddmut/)^99^. The surface electrostatic potential was evaluated using Adaptive Poisson-Boltzmann Solver (APBS) PyMOL plugin^100^. To further elucidate the impact of the R112H mutation, the interatomic interactions of wild type and AID^R112H^ were evaluated with Arpeggio^101^. Figures were prepared using PyMOL 3.1. The ssDNA (5’-AGTAGTTATCAGCTATGCTCAGTG-3’) was built using the program 3DNA^102^ and the AID models were generated using RoseTTAFold. The protein-DNA complex was formed by docking using HADDOCK software with default parameters^103^. The docked complex was selected from the top-ranked clustered structures.

### Statistics

Statistical analysis was performed as indicated in the figure legends. Statistics was performed using GraphPad Prism version 8. p≤ 0.05, significant; p>0.05, non-significant (ns).

## Acknowledgement

We are grateful to the staff KMA and the MTC animal facilities for their technical support. We thank Dr Javier di Noia (McGill University), Dr Kohsuke Imai (Tokyo Medical and Dental University), Dr Stephen Malin (Karolinska Institutet) for helpful discussions. This work was supported by postdoctoral fellowships from Olle Engqvist Byggmästare to M.H. and from Wenner-Gren Foundations to L.G.P., Karolinska Institutet PhD (KID) fellowships to R.D., E.D., T.Y., M.M.S.O., a PhD fellowship from Fundação para a Ciência e a Tecnologia and European Social Fund to M.M.S.O., a CAPES-STINT joint grant to R.G.G., V.C.d.A. and L.S.W., the Swedish Research Council and Cancer Society to L.S. and L.S.W., and Worldwide Cancer Research, Childhood Cancer Fund, Radiumhemmet Research Funds, and Karolinska Institutet to L.S.W. L.S.W. is a Ragnar Söderberg fellow in Medicine and holds a senior research position awarded by the Childhood Cancer fund. The CNIC is supported by the Instituto de Salud Carlos III (ISCIII), the Ministerio de Ciencia, Innovación y Universidades (MICIU) and the Pro CNIC Foundation, and is a Severo Ochoa Center of Excellence (grant CEX2020-001041-S funded by MICIU/AEI/10.13039/501100011033).

## Author Contribution

M.H. and L.S.W. designed the research, M.H., R.D., L.G.P., E.D., T.Y., R.C.V., M.M.S.O., R.G.G., S.S., C.S., N.V.K. and L.S.W. performed the experiments and analysed the data, D.K., M.A.Z, J.J.F., V.C.d.A., L.S., S.E.D. contributed with critical tools and discussions, M.H. and L.S.W. wrote the manuscript, and all authors edited the manuscript.

## Supplemental Figure legends

**Supplemental Figure 1.**
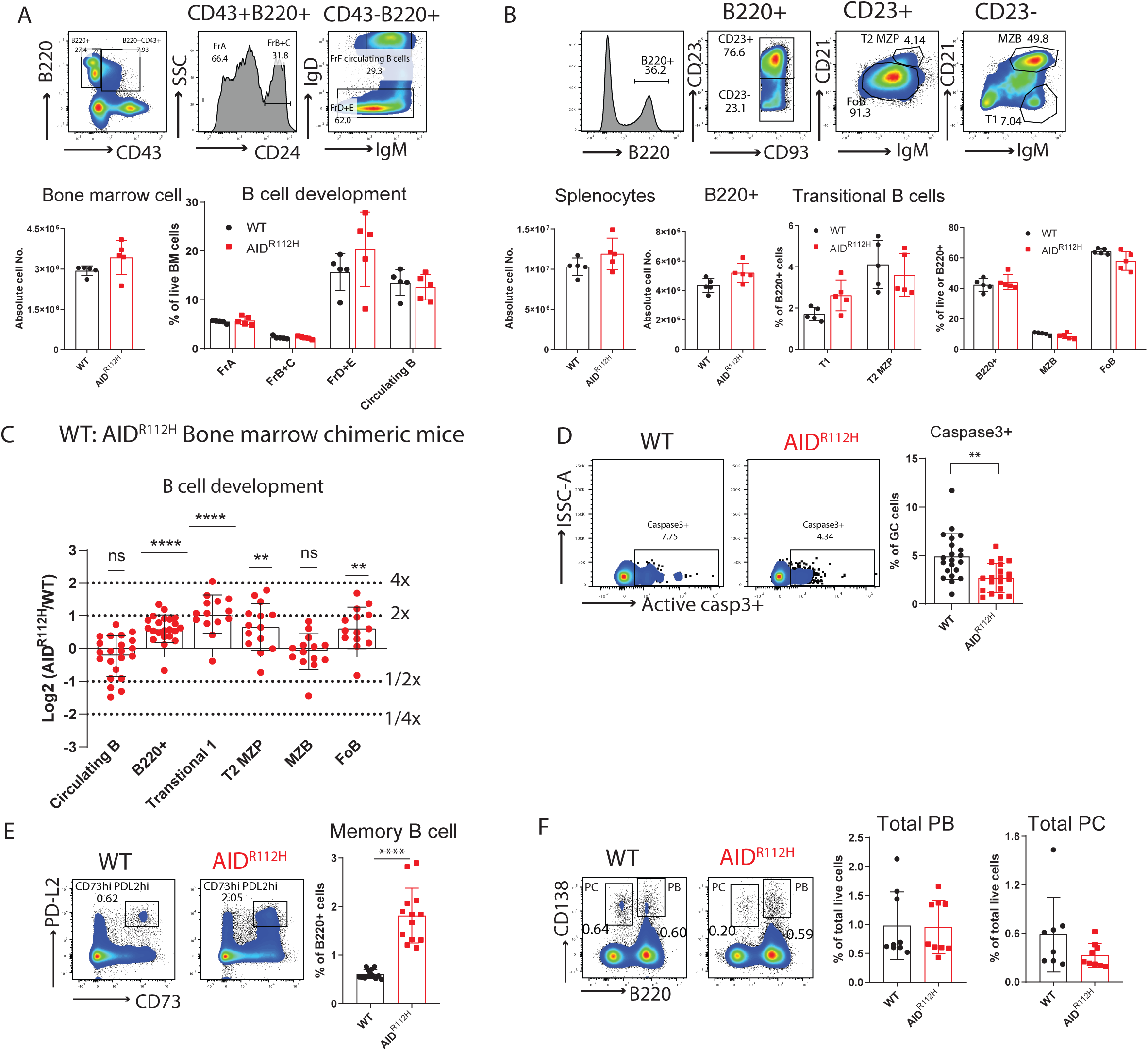
Antigen-independent B cell development in bone marrow and spleen. (A) B cell development in bone marrow by Hardy classification. Upper panels: representative FACS plots for gating. Lower panels: total cell number of bone marrow, p=0.032, WT vs AID^R112H^, unpaired *t* test (n= 5, one experiment); % of FrA (fraction A), FrB+C (fraction B+C), FrD+E (fraction D+E), non-significant when comparing WT (wildtype) vs AID^R112H^ in all groups. (B) B cell development in spleen. Upper panels: representative FACS plots for gating. Lower panels: quantifications, total spleen cells, non-significant; total B220^+^ cells, p=0.044, unpaired *t* test (n= 5, one experiment); % of T1 (Transitional 1), T2 MZP (transitional 2 marginal zone progenitor); B220^+^, MZB (marginal zone B cells), FoB (follicular B cells), non-significant for all groups, WT vs AID^R112H^, unpaired *t* test, (n= 5, one experiment). (C) B cell development in the bone marrow chimeric mice. y-axis: log2 of measured AID^R112H^/WT to grafted AID^R112H^/WT ratio, value 1=two-fold advantage, −1=two-fold disadvantage of AID^R112H^; B220+, Transitional 1: p<0.0001; Transitional 2 marginal zone progenitor, T2 MZP, Follicular B, FoB: p<0.01, one sample *t* test (n=14 to 22, three experiments). (D) Active Caspase3+ apoptotic cells in GC. FACS plots and quantification. WT vs AID^R112H^, p<0.01, *t* test (n=19-21, 4 experiments). (E) Memory B cell differentiation in spleen. FACS plot and quantification, p<0.01, unpaired *t* test, (n=13, three experiments). (F) Plasma cell differentiation in spleen. FACS plot and quantification: PB and PC, p>0.05. unpaired *t* test, (n=9, 2 experiments).

**Supplemental Figure 2.**
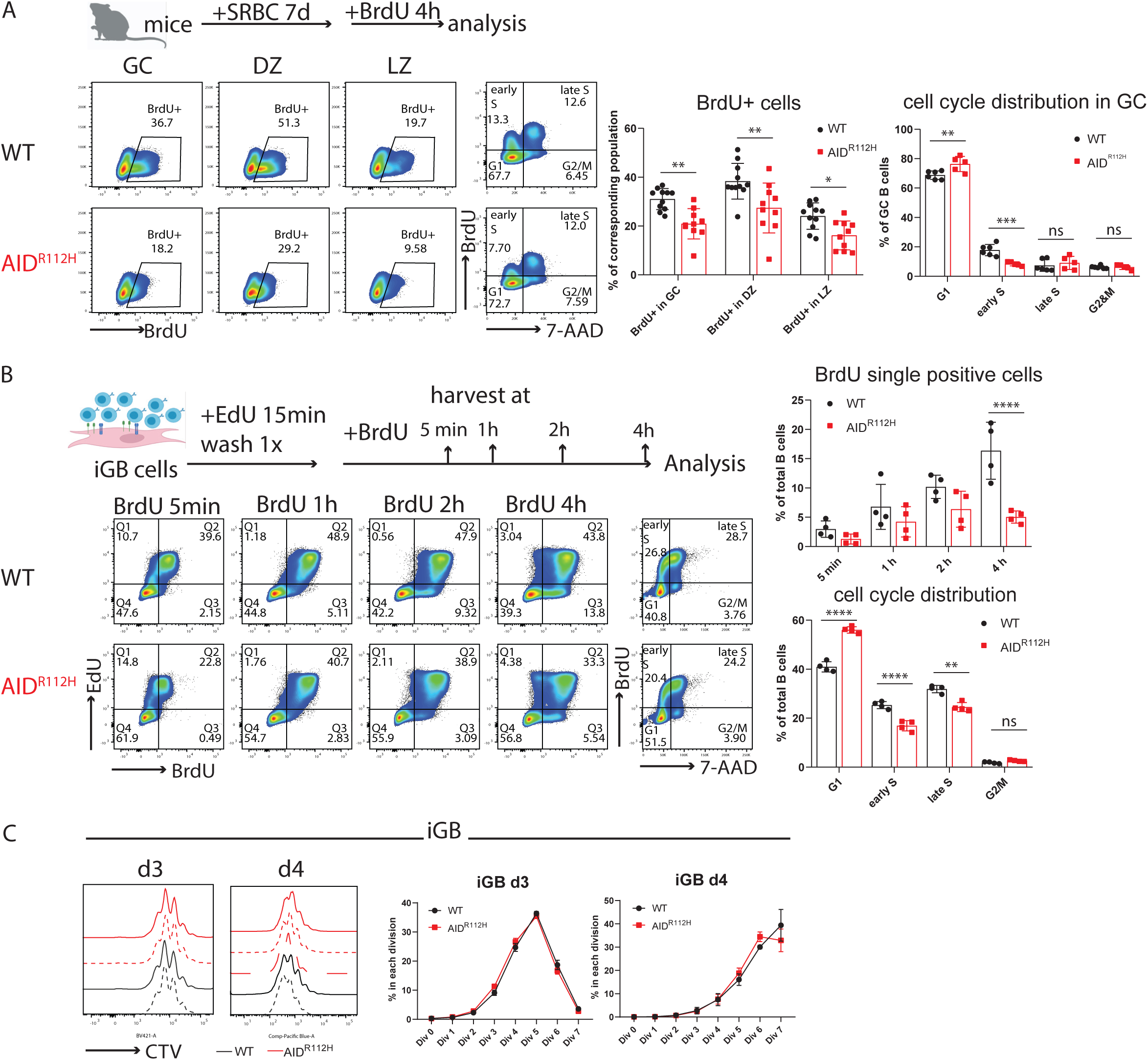
Reduced G1/S transition of AID^R112H^ iGB cells and GC B cells. (A) In vivo BrdU labeling of GC B cells on day 7 upon SRBC immunization. Schematics of experimental setup and representative FACS plots. Quantification: BrdU^+^ cells in GC: p= 0.0035; DZ: p=0.0014; LZ: p=0.0294; two-way ANOVA (n=10 to 11, 2 experiments). cell cycle distribution, G1: p=0.0027; early S: p=0.0002; two-way ANOVA (n=5 to 6, 1 experiment). BrdU mean fluorescence intensity (MFI) of BrdU^+^ cells in GC: p=0.078, unpaired t test, (n=5 to 6, 1 experiment). (B) EdU and BrdU double labeling in iGB cells. Schematics for experimental setup and representative FACS plots and Quantification, BrdU single positive cells at 4h: p<0.0001. Cell cycle distribution: G1, p<0.0001; early S, p<0.0001; late S, p<0.0001, (n=4, 2 experiments). (C) iGB cell proliferation rate measured by dilution of Cell Trace Violet (CTV). Left: histogram overlay of WT and AID^R112H^ cells on day 3 and day 4; Right: quantification of percentage of cells in each division. Value and error bar: mean±SD.

**Supplemental Figure 3.**
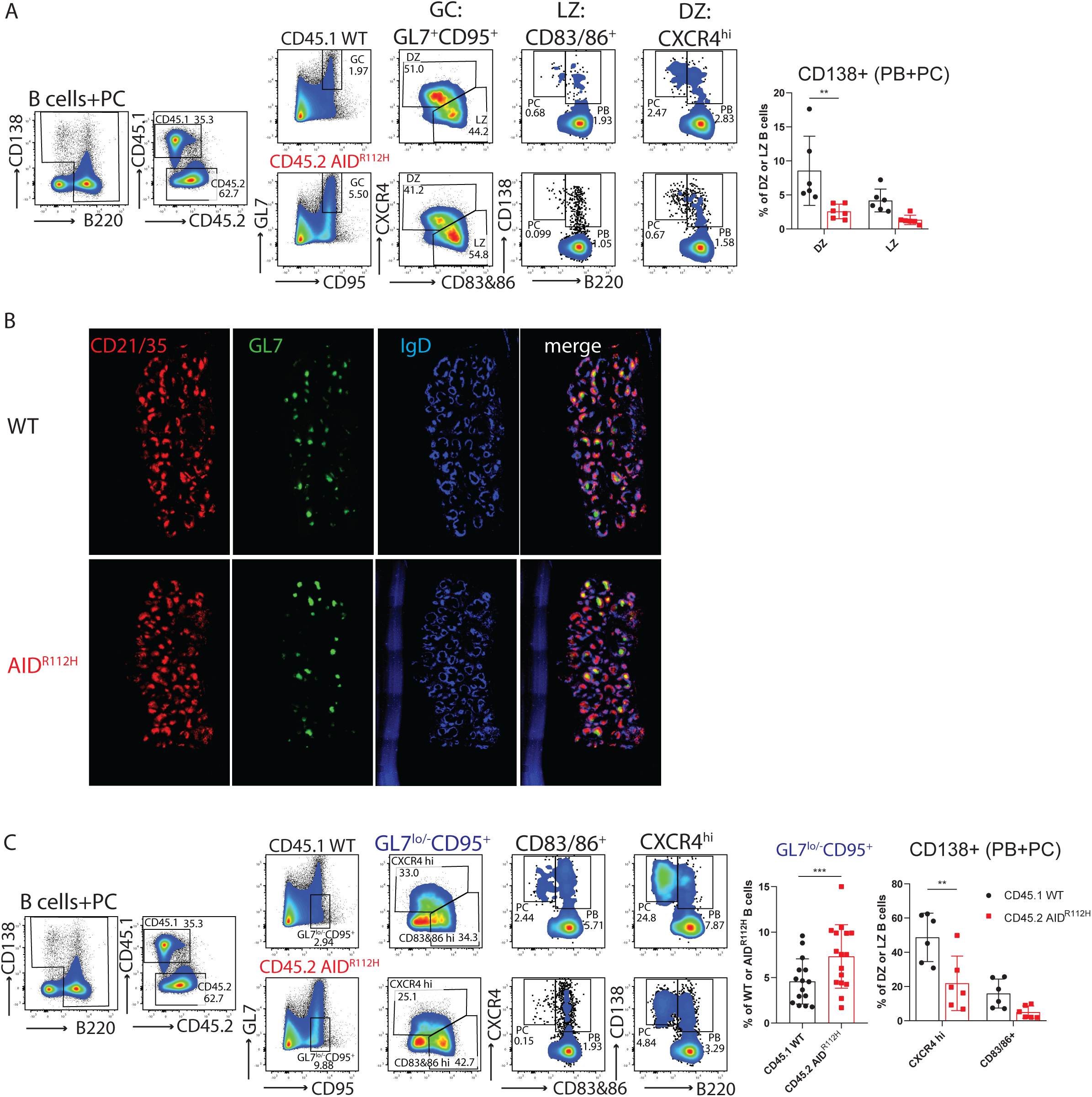
CD45.2 AID^R112H^ GC B cells have reduced plasma cell differentiation and accumulates GL7^lo/-^ tGC B cells in bone marrow chimeric mice. (A) Gating strategy to identify GC-derived PB and PC from CD45.1 WT cells and CD45.2 AID^R112H^ cells. PB: B220^+^CD138^+^, PC: B220^-^CD138^+^. Upper: representative FACS plots. Lower: quantification, CD138^+^ cells (PB+PC) in DZ and LZ. DZ: P=0.0024; LZ: non-significant. Two-way ANOVA, (n=6, 2 experiments); (B) Immunohistochemistry of GC from full spleen sections of WT and AID^R112H^ mouse 7 days after SRBC immunization. Red: CD21/35, Green: GL7, Blue: IgD. (C) Gating strategy of PB and PC in GL7^lo/-^CD95^+^ cell population. Upper: representative FACS plots; Lower: quantifications, GL7^lo/-^CD95^+^ cells in CD45.1 WT cells and CD45.2 AID^R112H^ cells, p<0.0001, paired *t* test, (n=15, 3 experiments). CD138^+^ cells in the different subsets from GL7^lo/-^CD95^+^ cell population: CXCR4^hi^: p=0.0014; CD83/86^+^: p= 0.225. two-way ANOVA, (n=6, 2 experiments).

**Supplemental Figure 4.**
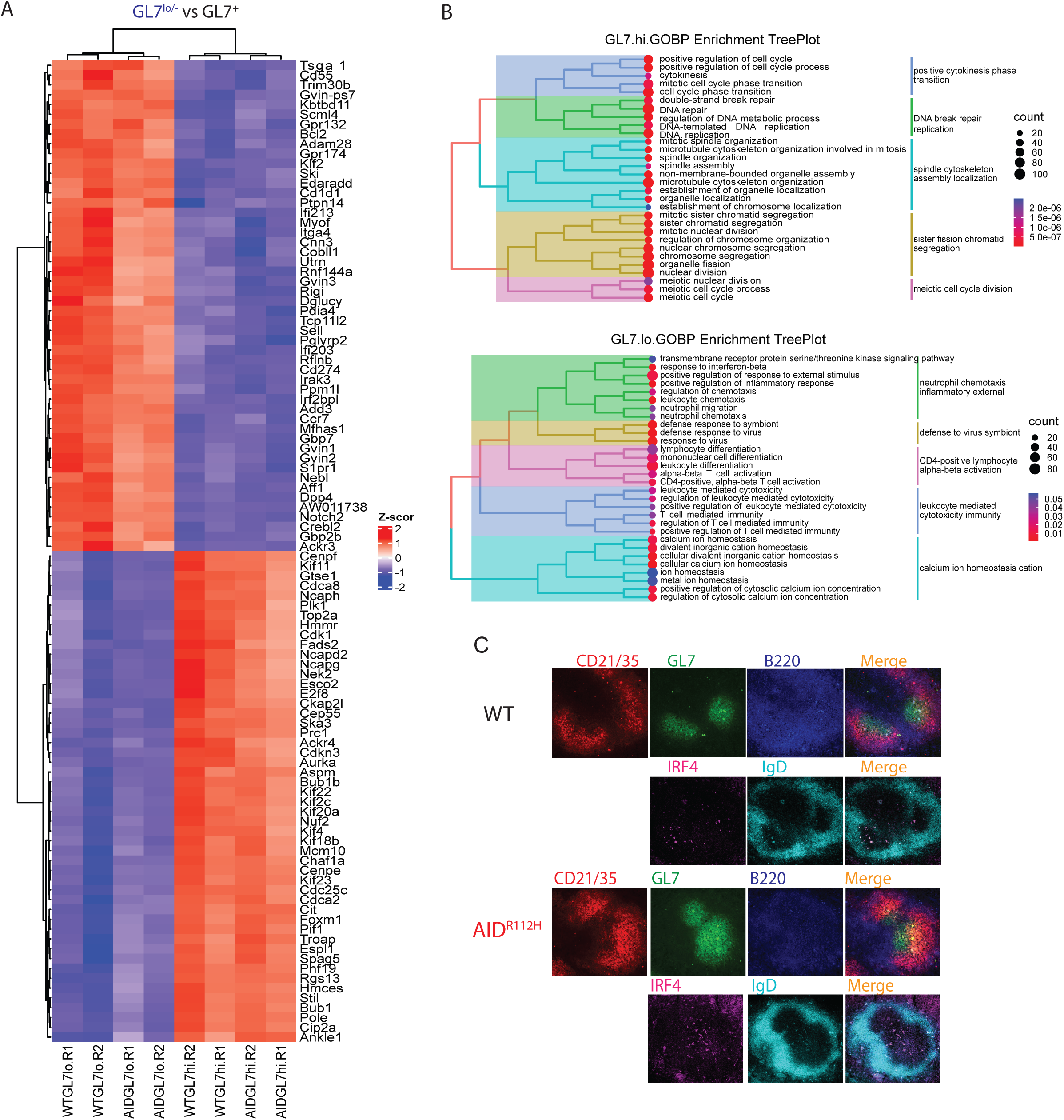
RNA sequencing data of GL7^lo/-^CD95^+^ versus GL7^+^CD95^+^cells sorted on day 7 after SRBC immunization. (A) RNA sequencing Heatmap for the top 50 differential expressed genes (DEGs) comparing GL7^lo/-^CD95^+^ vs GL7^+^CD95^+^. (B) GO term enrichment TreePlot for GL7^+^CD95^+^(GL7.hi.) cells and GL7^lo/-^CD95^+^(GL7.lo.) cells. (C) Immunohistochemistry to detect IRF4^hi^ cells within the GC area of WT mouse 7 days after SRBC immunization. Consecutive spleen sections with either CD21/35 (Red) + GL7 (Green) + B220 (Blue) or IRF4 (Purple) + IgD (Cyan).

**Supplemental Figure 5.**
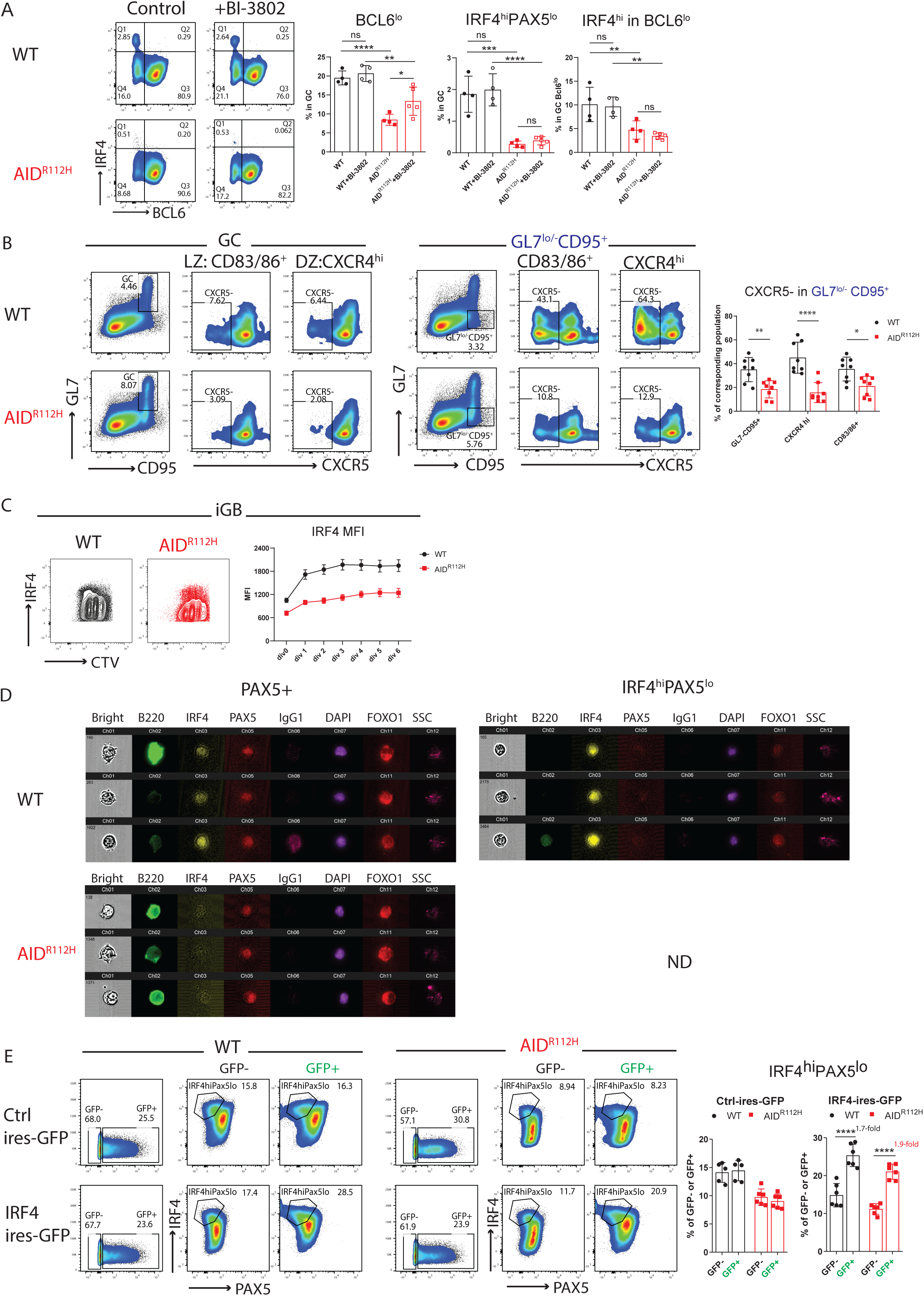
Impaired up-regulation of IRF4 is independent of BCL6 down-regulation and cell proliferation in GC. (A) Plasma cell differentiation in vivo during the GC response upon BI3802 treatment. Left: representative FACS plots. Right: quantification of BCL6^lo^: WT vs AID^R112H^, p<0.0001; WT+BI-3802 vs AID^R112H^+BI-3802: p<0.01; AID^R112H^ vs AID^R112H^+BI-3802: p<0.01; IRF4^hi^PAX5^lo^ cells: WT vs AID^R112H^, p<0.001; WT+BI-3802 vs AID^R112H^+BI-3802: p<0.0001; AID^R112H^ vs AID^R112H^+BI-3802: non-significant; IRF4^hi^ within the BCL6^lo^ population: WT vs AID^R112H^, p<0.01; WT+BI-3802 vs AID^R112H^+BI-3802: p<0.01; non-significant difference in treated vs non-treated, two-way ANOVA (n=4 to 5, one experiment). (B) Impaired CXCR5 down-regulation in AID^R112H^ tGC. FACS plots and quantification. GL7^lo^CD95^+^: p<0.01; CXCR4^hi^: p<0.0001; CD83/86+: p<0.05. one way ANOVA (n=8, 2 experiments). (C) Up-regulation of IRF4 versus cell division in iGB cells. Representative FACS plot and quantification, IRF4 Mean fluorescence intensity (MFI) vs cell division, data from 2 experiments. (D) Imagestream analysis of iGB cells on day 7. Representative images of PAX5^+^ and IRF4^hi^PAX5^lo^ cells from the iGB culture. Images were acquired with Imagestream flow cytometry (10000 cells acquired, 2 experiments), ND (non-detectable; no cells could be detected). (E) Overexpression of IRF4 in iGB cells by retroviral transfection. Left: representative FACS plots. Right: quantification of IRF4^hi^PAX5^lo^ cells in control and IRF4 transfected cultures. Ctrl-ires-GFP: no significant changes between WT GFP-vs WT GFP+; no significant changes between AID^R112H^ GFP-vs AID^R112H^ GFP+; IRF4-ires-GFP: p<0.0001, WT GFP-vs WT GFP+; p<0.0001, AID^R112H^ GFP-vs AID^R112H^ GFP+, two-way ANOVA (n=5 to 6, 2 experiments).

**Supplemental Figure 6.**
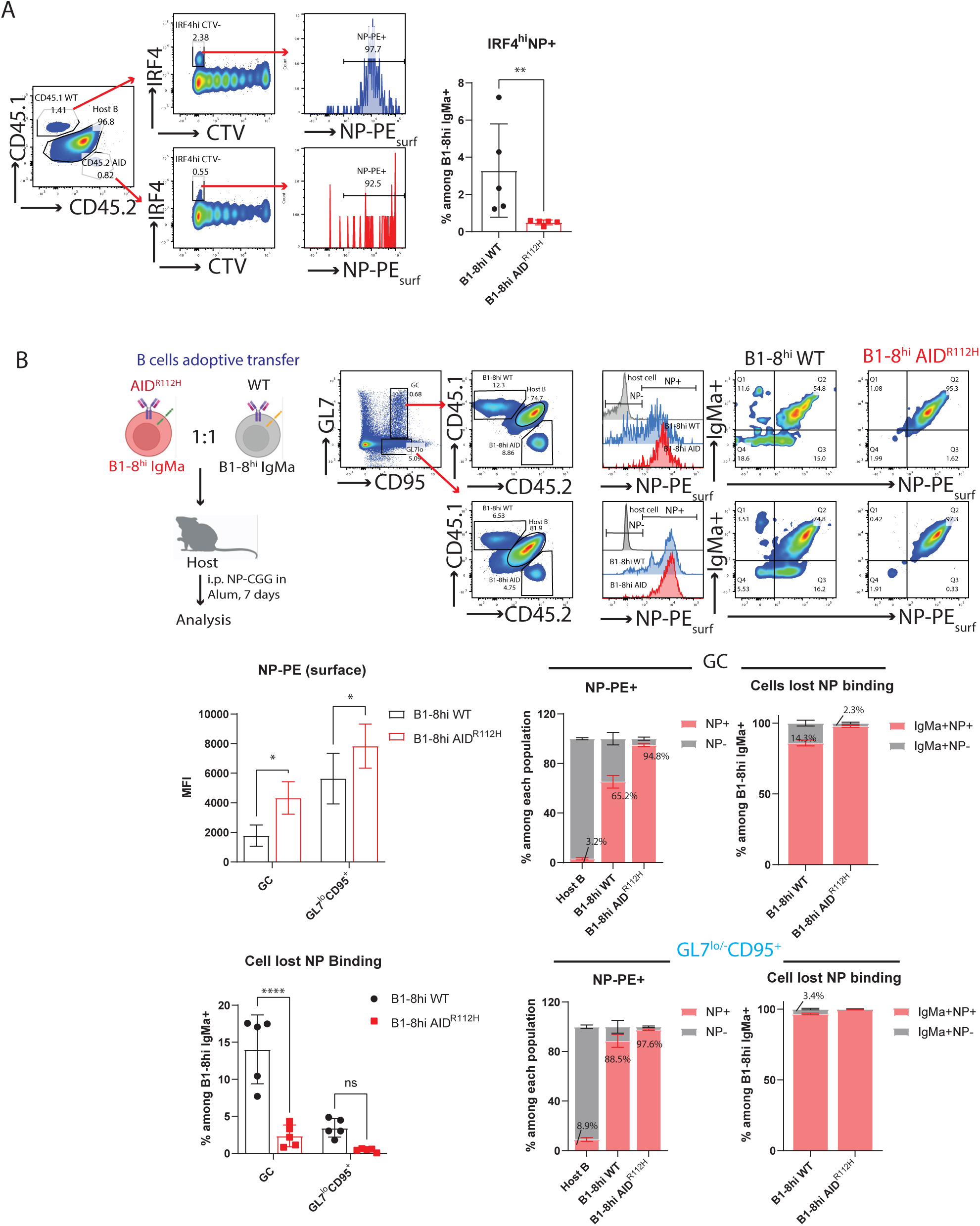
NP-specific plasma cell generation and affinity evolvement of B1-8^hi^ BCR in GC. (A) *In vivo* generation of IRF4^hi^NP^+^ cells from B1-8^hi^WT and B1-8^hi^AID cells. Left: gating strategy to detect NP-specific (surface staining) IRF4^hi^ cells by flow cytometry. Right: quantification, p<0.01, t test (n=5, 2 experiments). (B) NP binding capacity of B1-8^hi^ GC B cells. Upper: gating strategy to detect the NP binding capacity of B1-8^hi^ B cells in GC by FACS. Middle: left, surface NP-PE MFI in GC and GL7^lo^CD95^+^, B1-8^hi^WT vs B1-8^hi^AID cells: p<0.05, one way ANOVA (n=5, 2 experiments). Right: fraction of GC B cells that were NP-PE^+^ and NP-PE^-^ among the host B, B1-8^hi^WT and B1-8^hi^AID B cells; Fraction of BCR (IgMa)-expressing GC B cells that were NP-PE^+^ and NP-PE^-^ among B1-8^hi^WT and B1-8^hi^AID B cells. Lower: Left, quantification of IgM^+^ B1-8^hi^WT and B1-8^hi^AID cells that lost NP-binding in GC and GL7^lo^CD95^+^ cells, p<0.0001 for GC, non-significant for GL7^lo^CD95^+^, one way ANOVA (n=5, 2 experiments). Right: fraction of GL7^lo^CD95^+^ B cells that were NP-PE^+^ and NP-PE^-^ among the host B, B1-8^hi^WT and B1-8^hi^AID B cells; Fraction of BCR (IgM)-expressing GL7^lo^CD95^+^ B cells that were NP-PE^+^ and NP-PE^-^ among B1-8^hi^WT and B1-8^hi^AID B cells.

**Supplemental Figure 7.**
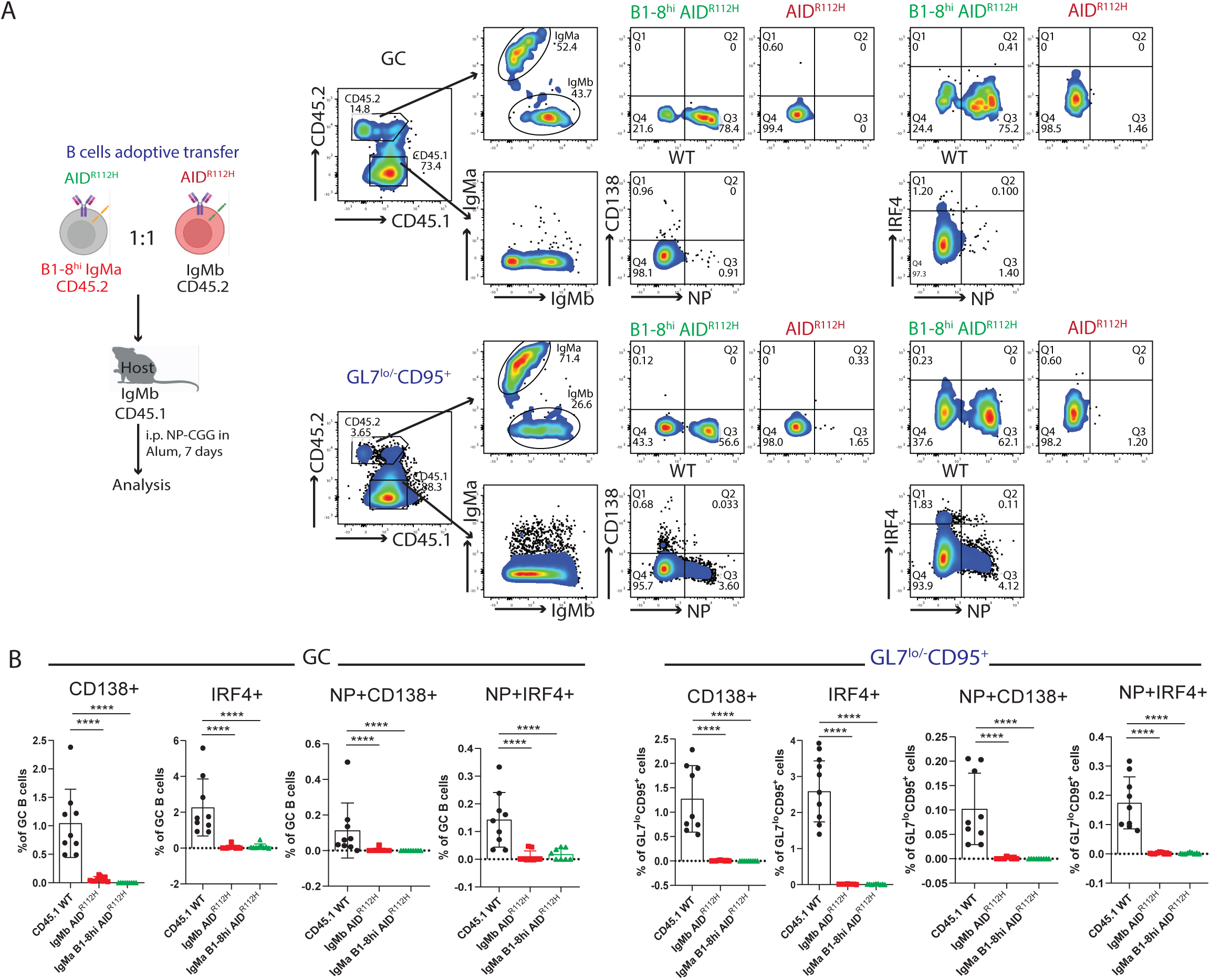
High affinity BCR is unable to rescue plasma cell differentiation of AID^R112H^ GC B cells. (A) Schematics of *in vivo* experimental setup. And gating strategy to analyze plasma cell differentiation capacity of B1-8^hi^AID^R112H^ (green), AID^R112H^ (red) and CD45.1 WT (black) cells. Representative FACS plots for GC B cells and GL7^lo/-^CD95^+^ B cells. (B) Quantification of CD138^+^, IRF4^+^, NP^+^CD138^+^ and NP^+^IRF4^+^ cells in GC and GL7^lo/-^ CD95^+^ cells. CD45.1 WT vs CD45.2 AID^R112H^: p<0.0001 for all groups. CD45.1 WT vs CD45.2 B1-8^hi^AID^R112H^: p<0.0001 for all groups. One-way ANOVA (n=9, 2 experiments). Value and error bar: mean±SD.

**Supplemental Figure 8.**
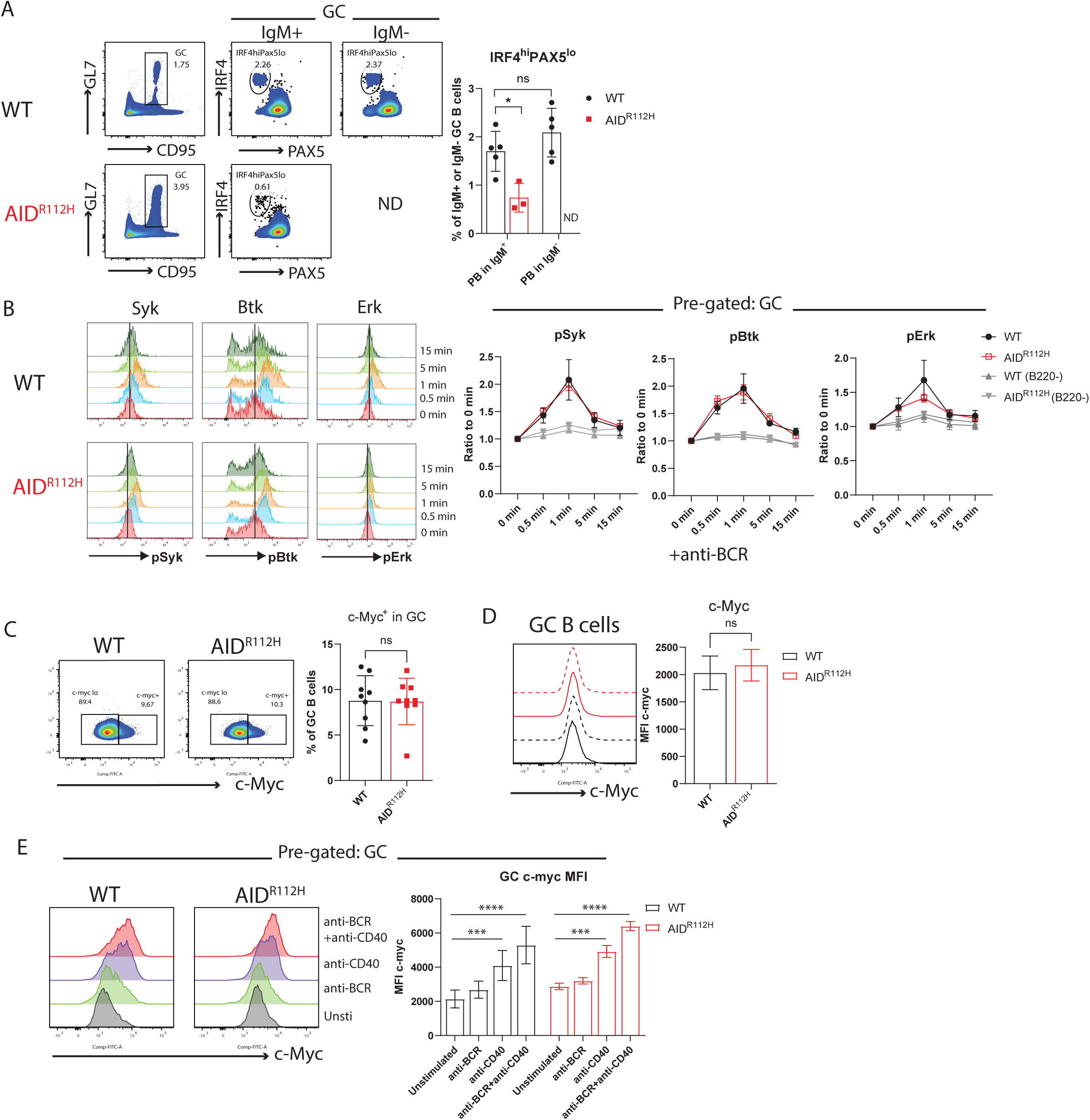
Ig class switching, BCR signaling and c-MYC expression in AID^R112H^ GC B cells. (A) Plasma cell differentiation capacity of switched (IgM-) and non-switched (IgM+) GC B cells. Left: Representative FACS plots. Right: quantification. WT vs AID^R112H^, PB in IgM+, p= 0.0102; IgM+ vs IgM-PB in WT GC, non-significant, unpaired t test (n=3-4, one experiment). (B) BCR signaling in WT and AID^R112H^ GC B cells. Left: FACS plots histogram overlay at different time points upon anti-BCR challenge. Right: quantification, WT vs AID^R112H^, non-significant, unpaired t test (n= 4, 2 experiments). (C) Fraction of c-MYC+ GC B cells. Left: representative FACS plots and gating strategy. Right: quantification, t test (n=9, 3 experiments). (D) c-MYC MFI in GC B cells. Right: histogram overlays to show WT (black, solid and dotted line) and AID^R112H^ (red, solid and dotted line) cells, t test (n=9, 3 experiments). (E) Up-regulation of c-MYC upon stimulation for 2 hrs in GC B cells. Left: histogram overlay of unstimulated and stimulated GC B cells. Right: quantification of c-MYC MFI, anti-BCR vs unstimulated, non-significant; anti-CD40 vs unstimulated, p<0.001 for both WT and AID^R112H^; anti-CD40+anti-BCR vs unstimulated, p<0.0001 for both WT and AID^R112H^; non-significant between WT and AID^R112H^ for corresponding stimulation conditions. Two-way ANOVA, (n=4 mice per condition, 2 experiments).

**Supplemental Figure 9.**
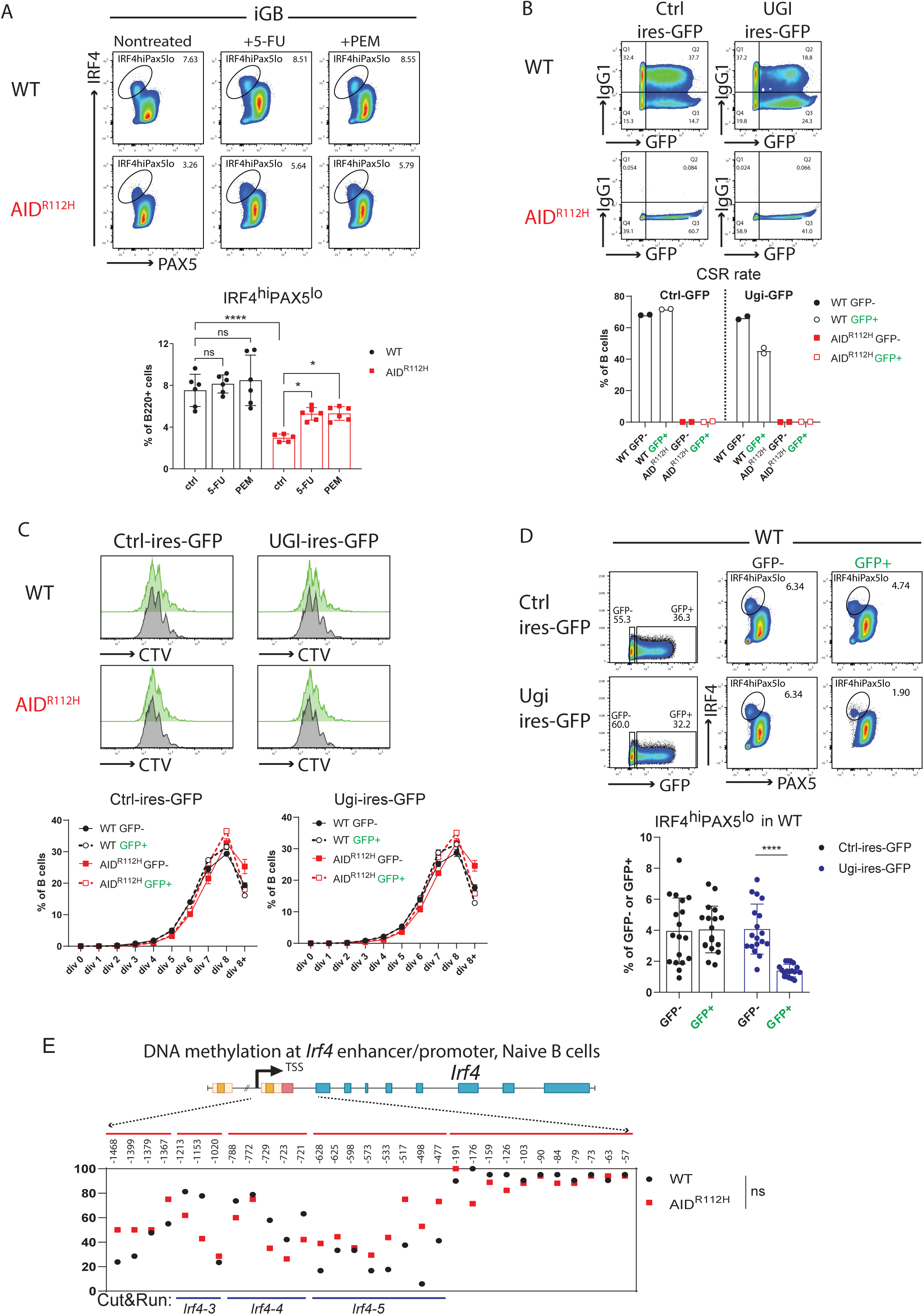
Impaired up-regulation of IRF4 independent of cell proliferation and response to stimulation. (A) Rescue of plasmablast differentiation upon 5-fluorouracil (5-FU) and pemetrexed (PEM) treatment in iGB culture. FACS plots and quantification. p<0.0001 WT vs AID^R112H^; p= 0.0133, AID^R112H^ vs AID^R112H^+5-FU; p=0.012, AID^R112H^ vs AID^R112H^+PEM, one-way ANOVA (n=6, 3 experiments). (B) Reduced Ig class switching to IgG1 upon UGI-ires-GFP retroviral transfection of iGB cells. Upper: representative FACS plots for control plasmid and UGI expressing plasmid transfection and Ig switching. Lower: quantification of Ig switching to IgG1 among the control plasmid and UGI plasmid transfected and non-transfected cells. (C) iGB Cell proliferation rate upon UNG inhibition by UGI. Upper: histogram overlay for non-transfected (gray) vs transfected (green) iGB cells (left: control plasmid, right: UGI plasmid). Lower: quantification of percentage of cells in each division in the transfected and non-transfected cells for both WT and AID^R112H^ iGB. Data from one representative experiment with replicates. (D) Inhibition of plasmablast differentiation by expressing Ugi peptide in WT iGB cells. FACS plots and quantification. Ctrl-ires-GFP: no changes between WT GFP-vs WT GFP+; Ugi-ires-GFP: p<0.0001 between WT GFP-vs WT GFP+, unpaired t test (n=18, 2 experiments). (E) CpG DNA methylation status at the *Irf4* locus in naïve B cells from WT and AID^R112H^ mice. Top: schematics for the *Irf4* gene locus and the region examined for DNA methylation using bisulfite sequencing. DNA methylation level at CpG sites, number above the graph shows the CpG position relative to first exon. For each CpG site, 15-30 clones were analyzed, data from 2 independent experiments.

**Supplemental Figure 10.**
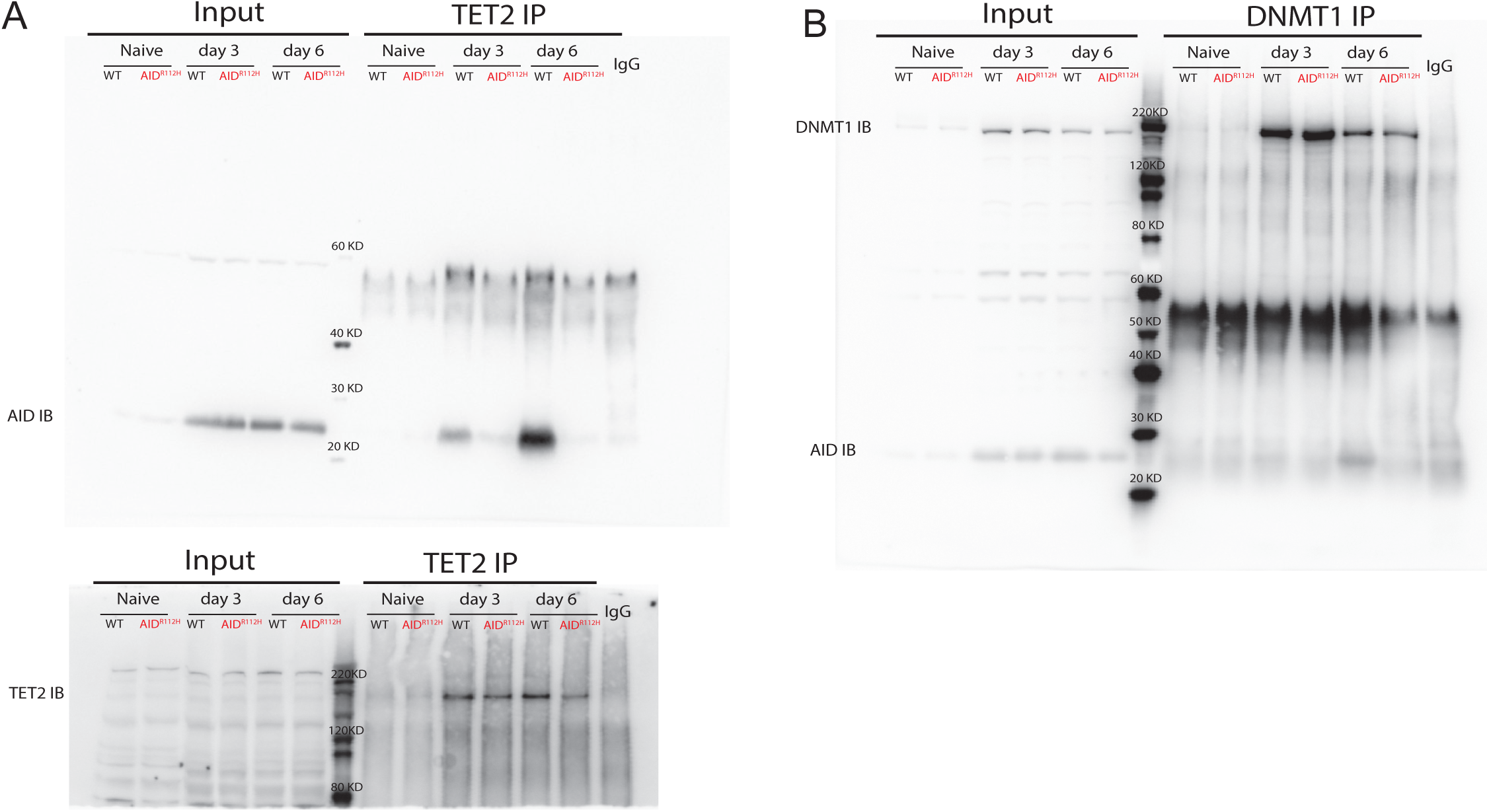
AID interacting proteins and chromatin association in iGB cells. (A) TET2 Immunoprecipitation on day 3 and day 6. Immunoblot for AID (ActM, upper panel) and TET2 (Abcam, lower panel). (B) DNMT1Immunoprecipitation on day 3 and day 6. Immunoblot for AID (ActM) and DNMT1 (Merck).

**Supplemental Figure 11.**
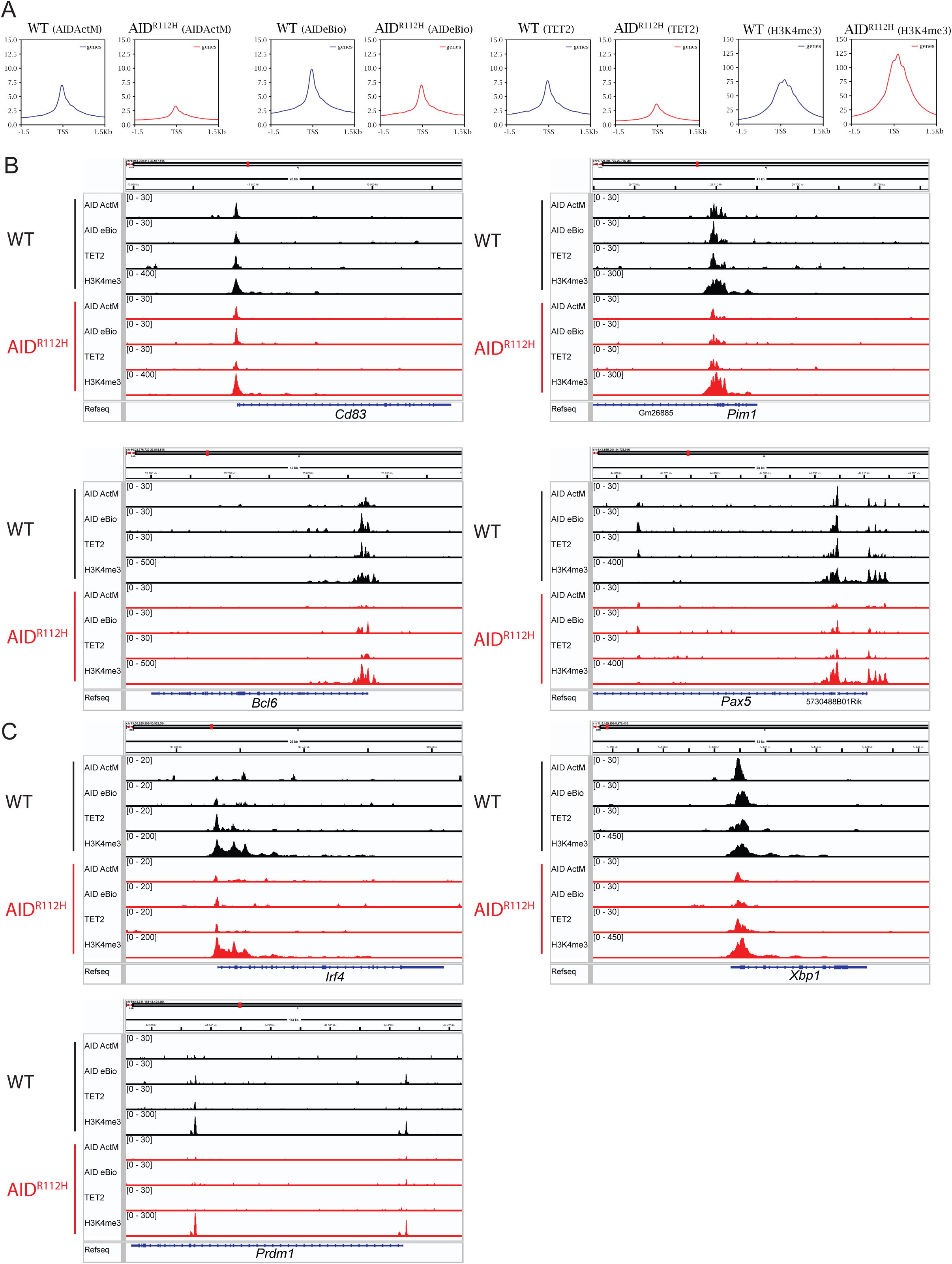
Association of AID, TET2 and histone mark H3K4me3 to AID off-target genes. (A) Cut&Run sequencing to detect global chromatin association of AID, TET2, H3K4me3 in iGB cells on day3. Upper: average enrichment profiles near TSS (−1.5kb to +1.5kb). (B-C) Cut&Run-sequencing Peak profiles of genes involved in the GC response including *Cd83, Pim1, Bcl6* and *Pax5* (B) and for genes involved in plasma cell differentiation including *Irf4, Xbp1* and *Prdm1*(C). The enrichment profiles represent the mean of two separate experiments sequenced at the same time.

**Supplemental Figure 12.**
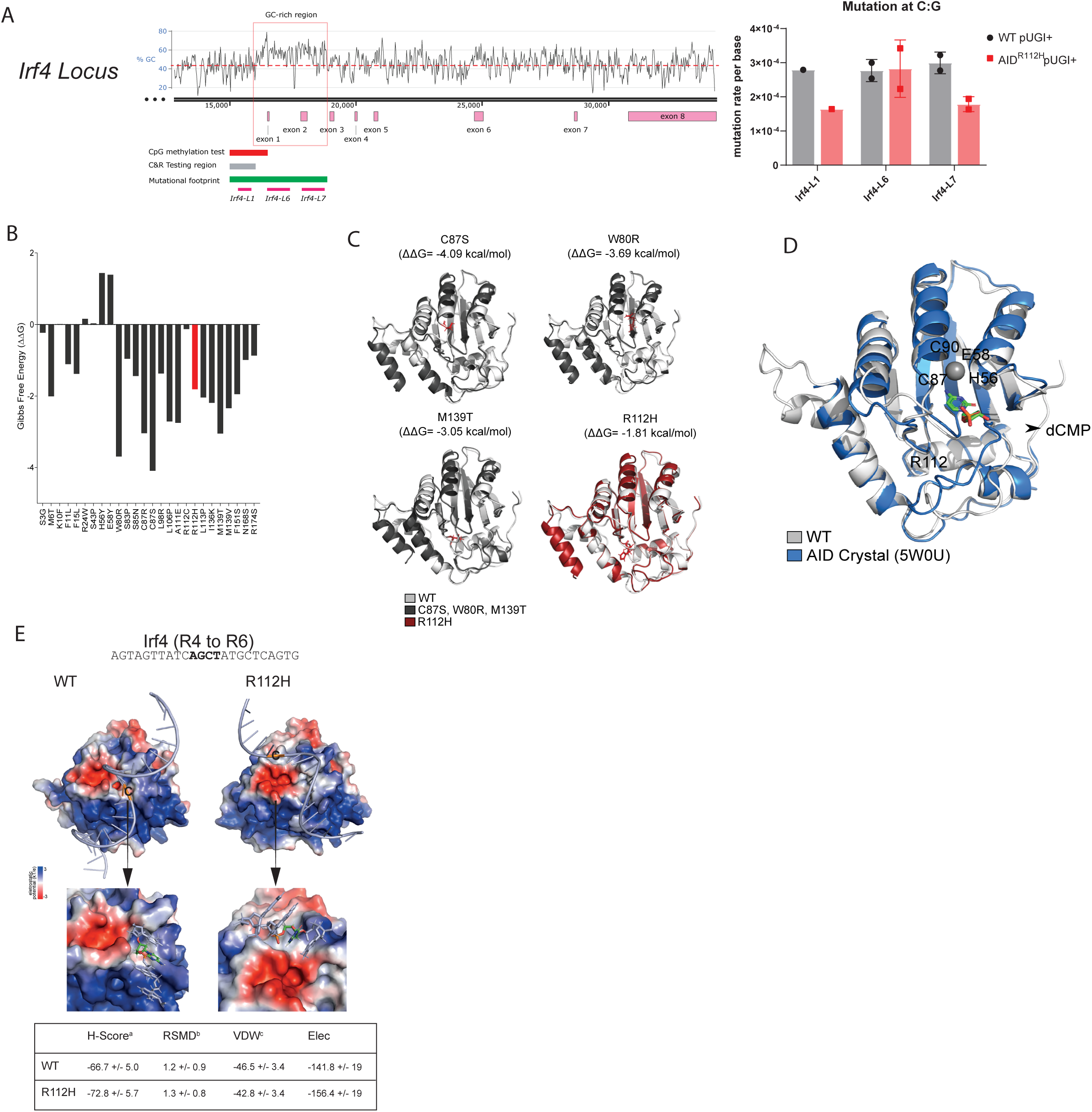
AID-dependent mutational footprint at the *Irf4* locus and structure analysis of AID mutations including R112H. (A) Mutational footprint of AID at the *Irf4* locus. Left: GC-content analysis of *Irf4* gene locus, GC-rich region highlighted in red square. Positions of exons, region for demethylation test, Cut&Run test and mutation footprint are indicated below. Right: Quantification of C to T and G to A mutation rate at the indicated regions. (B) DDMut scores of selected mutations in AID. Red marks R112H mutation, black marks other AID mutations. (C) The 3D models of wild type and mutated AID (predicted) aligned together. (D) Structure of AID from RoseTTAFold aligned to AID crystal bound to dCMP (PDB:5W0U). Catalytic residues H56, E58, C87, C90, and the residue R112 highlighted. (E) *Irf4* probe binding to AID active-site cleft. Left upper: *Irf4* probe binding to WT and R112H AID protein. Left lower: Zoomed views of the simulated docked complex. AGCT motif is displayed as licorice representation, cytosine is colored in green (carbon atoms). Right: The values for the top ranked structures. The values were obtained from HADDOCK software. a HADDOCK score. b RMSD (Å) value from the overall lowest-energy structure. c Van der Waals energy (kcal/mol). d Electrostatic energy (kcal/mol).

---

**Supplemental Table I.**
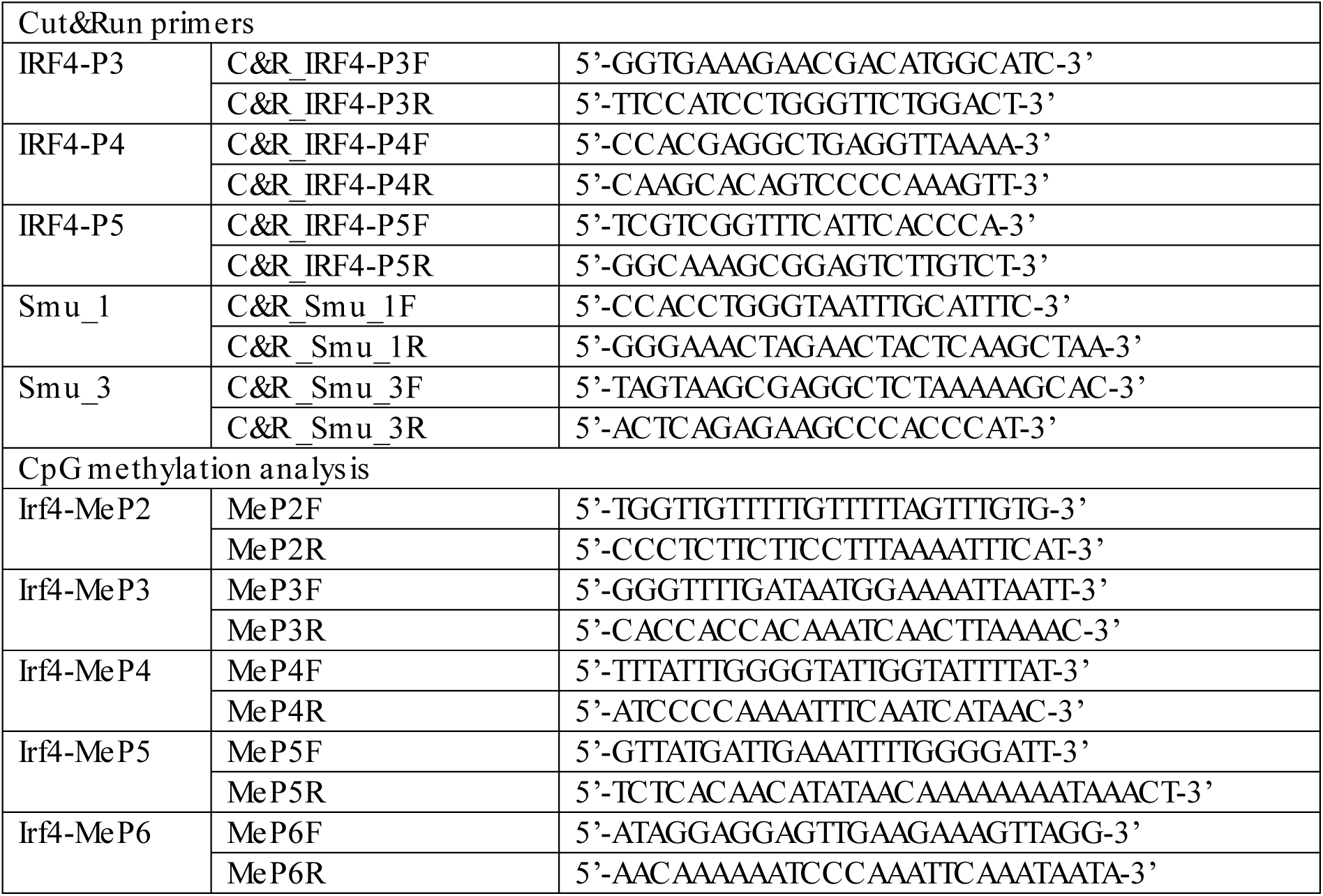
Primer sequences.

